# The Colon Mucosal Sialylglycome Is Redox-Regulated by the Golgi Enzyme QSOX1

**DOI:** 10.1101/2022.05.03.490496

**Authors:** Tal Ilani, Nava Reznik, Noa Yeshaya, Tal Feldman, Patrick Vilela, Zipora Lansky, Gabriel Javitt, Michal Shemesh, Ori Brenner, Yoav Elkis, Neta Varsano, Nathan Murray, Parastoo Azadi, Ana M. Jaramillo, Christopher M. Evans, Deborah Fass

## Abstract

Mucus shields the intestinal epithelium from pathogens and provides a supportive environment for commensal bacteria. Mucus is composed of enormous, heavily glycosylated proteins called mucins, which become disulfide crosslinked in a multi-step biosynthetic pathway culminating in the Golgi apparatus and secretory granules of goblet cells. We observed that knockout mice lacking the Golgi-localized disulfide catalyst QSOX1 produced poorly protective colon mucus, were hypersensitive to induced colitis, and had an altered microbiome. The initial hypothesis arising from these observations was that QSOX1 catalyzes disulfide crosslinking of mucins. Contrary to this hypothesis, the disulfide-mediated polymerization of mucins and related glycoproteins proceeded normally without QSOX1. Instead, we found that QSOX1 forms regulatory disulfides in Golgi glycosyltransferases and thereby promotes effective sialylation of the colon glycome. Our findings reveal that enzymatic control of Golgi redox state impacts glycan elaboration in goblet cells, and that this pathway is crucial for maintaining mucosal function.

---

Gastrointestinal health depends on the successful construction and performance of protective mucus secreted by goblet cells into the gut lumen^1, 2^. Mucus is a hydrogel, composed of networks of mucin glycoproteins, that serves as a physical barrier separating pathogenic microorganisms, viruses, and parasites from the underlying epithelium^3, 4^. Mucins also feed the symbiotic gut microbiome^5^ and interact with proteins that aid in tissue regeneration and repair^6^. Some progress has been made in understanding the molecular structures of mucins^7–10^, but factors that contribute to mucin functional organization and maintenance in vivo are poorly understood. The mucosal barrier defects associated with inflammatory bowel diseases have been suggested to arise post-transcriptionally^11^, e.g., at the levels of protein synthesis, oligomerization, and post-translational modification. An improved mechanistic understanding of mucus construction in goblet cells may offer new opportunities to strengthen possible weak links in the mucin scaffold and enhance the protective activity of the hydrogel, thereby improving resistance to infections and inflammation^12^.

Mucin glycoproteins form long polymers linked by disulfide bonds acquired in a step-wise intracellular assembly process. Intermolecular disulfide bonds crosslink two mucin carboxy termini to form dimers in the endoplasmic reticulum (ER)^13^. Subsequently, multiple mucin dimers are connected to form polymers by disulfide bonding of their amino termini in the Golgi apparatus or other post-ER compartments^14, 15^. The carboxy-terminal crosslinking^16^, along with extensive intramolecular disulfide bonding, is likely to be carried out by the numerous thiol-disulfide oxidoreductases that comprise the general oxidative protein folding machinery of the ER^17^, with the aid of specific factors such as Agr2^18, 19^. In contrast, the potential players in disulfide-mediated mucin polymerization in the Golgi are more limited. Quiescin sulfhydryl oxidase 1 (QSOX1), a catalyst of disulfide bond formation localized to the Golgi in various cell types^20^, is expressed at high levels in intestinal goblet cells^21^, where mucus is produced. It was thus reasonable to speculate that QSOX1 catalyzes the second step of intermolecular disulfide bonding in mucin polymer biosynthesis. In this study, we set out to test this hypothesis.

Assays of mucin bioassembly are complicated by the same features that give these proteins their extraordinary structural and mechanical capabilities. Mucins stand out for their unusual lengths, extensive glycosylation, and large number of intramolecular disulfide bonds. For example, human intestinal mucin 2 (MUC2) is more than 5000 amino acids long, has over 100 disulfides, and can bear up to five times its protein weight in O-linked glycans^22, 23^. Furthermore, goblet cells are highly differentiated factories for mucin production^24^ that are not easily adapted to cell culture and genetic manipulation, though airway epithelial cell lines have been used successfully to study mucin mesoscale structure^25^. To ensure a physiological context for studying the role of QSOX1 in mucin assembly, we generated and examined QSOX1 knockout (KO) mice, focusing our analysis on the colonic mucosa. Our findings show that QSOX1 activity is indeed related to mucus hydrogel production, but that Golgi disulfide bonding impacts mucus quality and functionality by an unanticipated pathway.

## Results

### QSOX1 localizes to the Golgi apparatus of colon epithelial cells

QSOX1 is expressed at high levels during development^26, 27^ and in a cell-type specific manner in particular adult tissues^21^. Single-cell transcriptomics data from large intestine reported by others^28^ and confirmed by us show that QSOX1 levels are highest in goblet cells. Consistent with QSOX1 intracellular localization in other cell types, QSOX1 immunofluorescence co-localized with Golgi marker GM130 in cells isolated from murine colon epithelium (Fig. 1a).

**Fig. 1.**
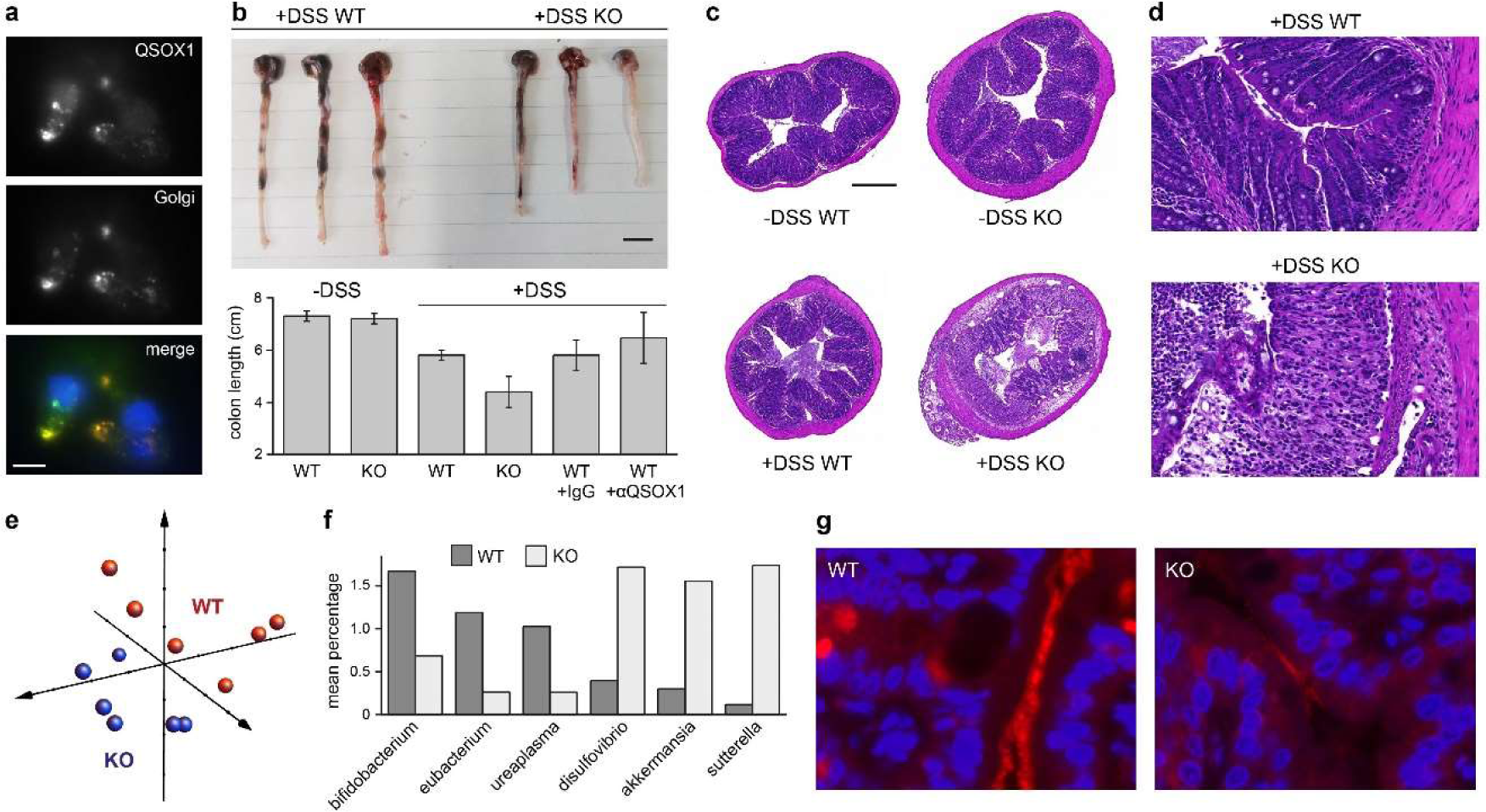
QSOX1 KO mice are hypersensitive to induced colitis and show alterations in their microbiome. **a,** Immunofluorescence labeling of QSOX1 and a Golgi marker (GM130) in epithelial cells isolated from murine colon. QSOX1 is green, GM130 is red, and DAPI staining of nuclei is blue in the merged image. Scale bar is 10 µm. **b,** Left, average colon length of WT and QSOX1 KO mice following indicated treatments. (n ≥ 3). αQSOX1 is MAb316.1, which inhibits murine QSOX1^33^. Right, representative images of colons following DSS treatment. Scale bar is 1 cm. **c,** Colon cross-sections stained with H&E show severe damage to the epithelium in DSS-treated KO mice. Scale bar is 500 μm. d, Damage to epithelium and immune cell infiltration in DSS-treated QSOX1 KO colons. Scale bar is 50 μm. e, Principal coordinate plot of weighted Unifrac data from amplicon sequencing of feces from WT and KO mice using 16S universal eubacterial primers. The three vectors presented exhibit more than 70% of the variation among the groups. f, Mean percentage is displayed for genera that differ significantly between WT and KO fecal samples and that represent at least 1% of the sequences for either genotype. g, Fluorescence in situ hybridization of 16S rRNA (red). Blue indicates DAPI staining of nuclei. Scale bar is 10 µm.

### QSOX1 knockout mice are highly susceptible to induced colitis and have an altered microbiome

QSOX1 KO mice were generated using commercial embryonic stem cells in a C57BL/6 background. KO mice were viable, fertile, had a normal lifespan, and showed no overt physical or behavioral abnormalities, in accordance with a previous report^29^. In particular, QSOX1 KO mice did not display the growth retardation, occult blood loss, and rectal prolapse exhibited by mice lacking Muc2, the major intestinal gel-forming mucin^30^. To determine whether QSOX1 nevertheless contributes to the physiology of the intestinal mucosa, we challenged the gastrointestinal tract. KO and control wild-type (WT) littermates were treated with dextran sodium sulfate (DSS) via their drinking water according to an established protocol for inducing acute colitis^31, 32^. QSOX1 KO mice were found to be more sensitive, displaying severe disease symptoms at day four to five when WT littermates were only mildly affected (clinical scores were 1-2 for WT and 3-4 for KO, see Methods). Without DSS treatment, QSOX1 KO mice had similar colon lengths and crypt organization as WT (Fig. 1b). After DSS treatment, however, colons of KO mice were shorter than those of WT, indicating more widespread epithelial loss. Administration of QSOX1 inhibitory antibodies^33, 34^ did not enhance sensitivity of WT colons to DSS, the significance of which will be discussed. Colons of DSS-treated KO mice showed pervasive disruption of ultrastructure, while colons of WT mice were largely unaffected (Fig. 1c). Additionally, massive immune cell infiltration was seen in KO colons but not in WT colons after DSS treatment (Fig. 1d).

Though untreated WT and QSOX1 KO animals did not show appreciable differences in colon ultrastructure, marked differences in microbiome composition were detected. Amplicon sequencing of bacterial 16S RNA from colon feces of co-housed WT and QSOX1 KO mice revealed clear clustering of genera according to mouse genotype (Fig. 1e). The specific bacterial genera with relative populations elevated in the KO mice were Desulfovibrio and Sutterella, previously identified as associated with active colitis^35^, as well as Akkermansia, which subsists on mucins^36^ and is considered a beneficial probiotic^37^ (Fig. 1f and Extended Data Fig. 1). The genera Bifidobacterium, Eubacterium, and Ureaplasma were elevated in WT mice compared to the KO (Fig. 1f and Extended Data Fig. 1). The wide range of genera found to be significantly different between WT and KO feces samples shows that lack of QSOX1 impacts the colon microbiome even in the absence of additional stress or challenge. Interestingly, fluorescence in situ hybridization to detect 16S rRNA and reveal the spatial distribution of bacteria in the gut showed weak signal in KO compared to WT colon sections (Fig. 1g). Experiments described below will illuminate the cause of this difference.

### QSOX1 knockout mice have defects in colon mucus

As mucosal barrier function and bacterial colonization depend on colon mucus quantity and quality, we directly analyzed colon mucins in the QSOX1 KO mice. Transcript levels of mucin genes and factors required for mucin production and trafficking were increased in QSOX1 KO colons compared to WT colons (Extended Data Fig. 2). However, an inspection on the protein level revealed decreased Muc2 in the luminal spaces of QSOX1 KO colon cross-sections. Lower luminal Muc2 immunofluorescence labeling was reproducible with multiple antibodies recognizing either the Muc2 amino terminus (Fig. 2a,b) or carboxy terminus (Fig. 2c). Most strikingly, the colons of WT mice showed a continuous, sharp line of Muc2 coating the epithelial layer, whereas the KO mice showed only diffuse and weaker extracellular Muc2 labeling (Fig. 2d). A complementary perspective was obtained by a longitudinal colon cut to expose the lumen and visualize the secreted mucus from above. Examined either by immunofluorescence (Fig. 2e and Extended Data Fig. 3) or scanning electron microscopy (SEM) (Fig. 2f), the lumen of WT mice showed mucin bundles covering the epithelium, contrasting with either naked, exposed cell surface or small, disorganized mucin patches in the KO. The lack of luminal mucus in fixed ex vivo colon samples from QSOX1 KO mice is consistent with the low 16S RNA labeling (Fig. 1g), as bacteria inhabit colon mucus.

**Fig. 2.**
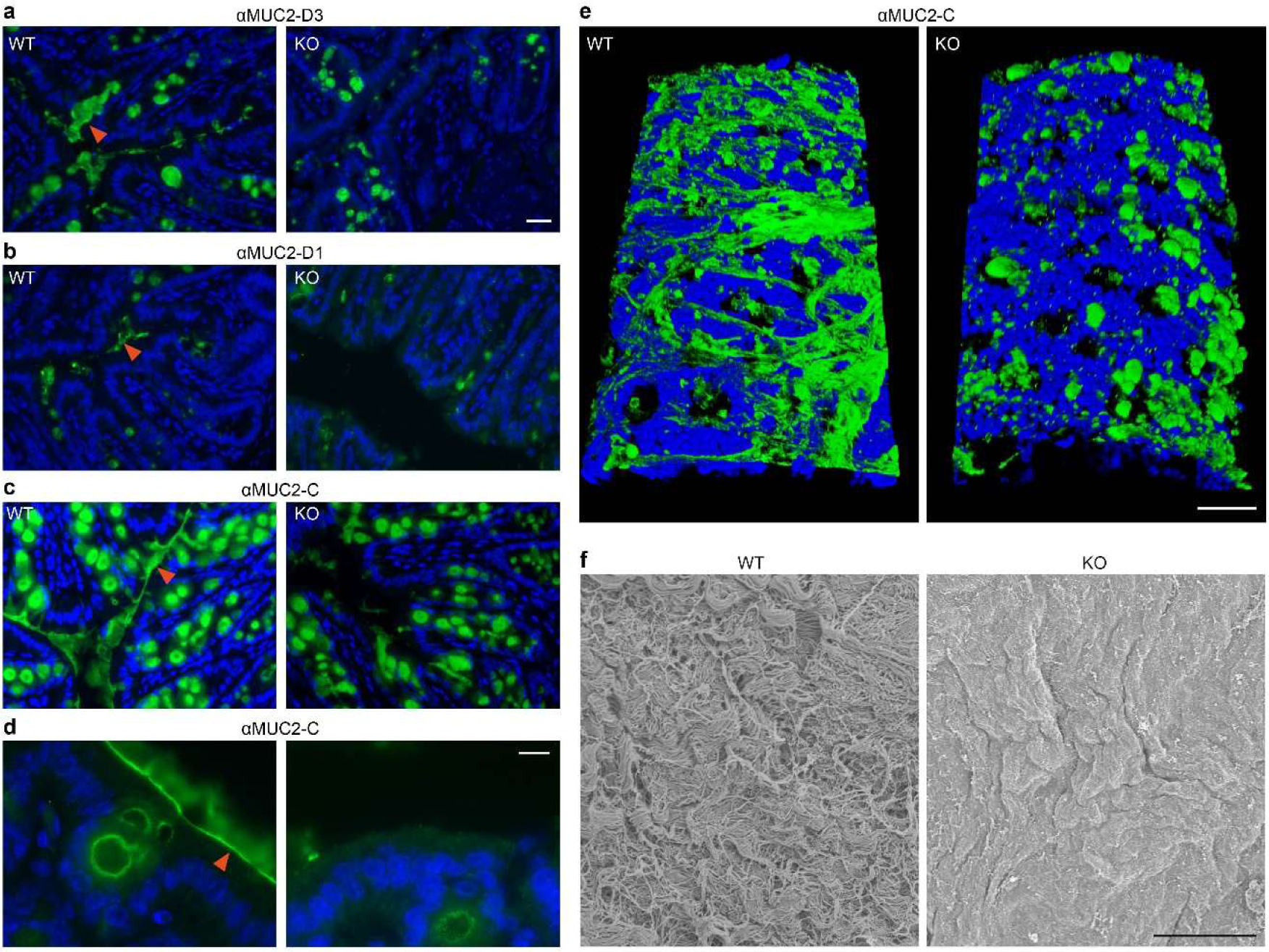
Perturbed colon mucins in QSOX1 KO mice. **a,** Cross-sections of WT and KO colons immunolabeled (green) for Muc2 with an antibody recognizing the D3 assembly in the amino terminal segmeent (αMuc2-D3)^8^. Blue is DAPI staining of nuclei. Orange arrowhead indicates secreted mucin. Scale bar is 20 µm and applies also to panels b and c. **b,** As in panel a except using an antibody recognizing the D1 assembly in the amino terminal segment (αMuc2-D1). The antibody was raised against the human ortholog but is cross-reactive with murine Muc2. **c,** As in panel a except using an antibody recognizing the Muc2 carboxy terminus (αMuc2-C). **d,** A higher magnification image of a WT colon cross-section shows a sharp line of Muc2 coating the epithelium (orange arrowhead), covered by diffuse mucin labeling. These features are absent from KO colons. Scale bar is 10 µm. **e,** Immunolabeling (αMuc2-C; green) of the lumen of longitudinally cut colon sections. Blue is DAPI staining of nuclei. Scale bar is 50 µm. Incomplete coverage of the colon lumen by mucin strands even in WT may be due to limitations of the fixation procedure. f, SEM micrographs reveal thick mucus coating the WT colon in contrast to the exposed epithelial cell surface of the KO colon. Scale bar is 5 µm.

The poor properties of QSOX1 KO colon mucus raised the question of where in the process of mucin production, assembly, packaging, and secretion the molecular defects arise. If QSOX1 activity is required in the Golgi or during goblet cell granule loading, then the packing of granules might be affected by the lack of QSOX1. To examine goblet cell secretory granules at high resolution, transmission electron microscopy (TEM) was performed on thin colon sections. In both WT and KO animals, full mucin granules were observed (Fig. 3a). Many granules in both WT and KO mice had well-delineated, polygonal sub-compartments, but there seemed to be an increased number of immature granules with rounder sub-compartments in the KO, consistent with increased mucin production and turnover rates. Together these observations establish that QSOX1 KO mice synthesize and package mucins. Upon release, however, the defect becomes clear, with the KO mice being unable to produce or retain a densely assembled mucin network (Fig. 2a-e).

**Fig. 3.**
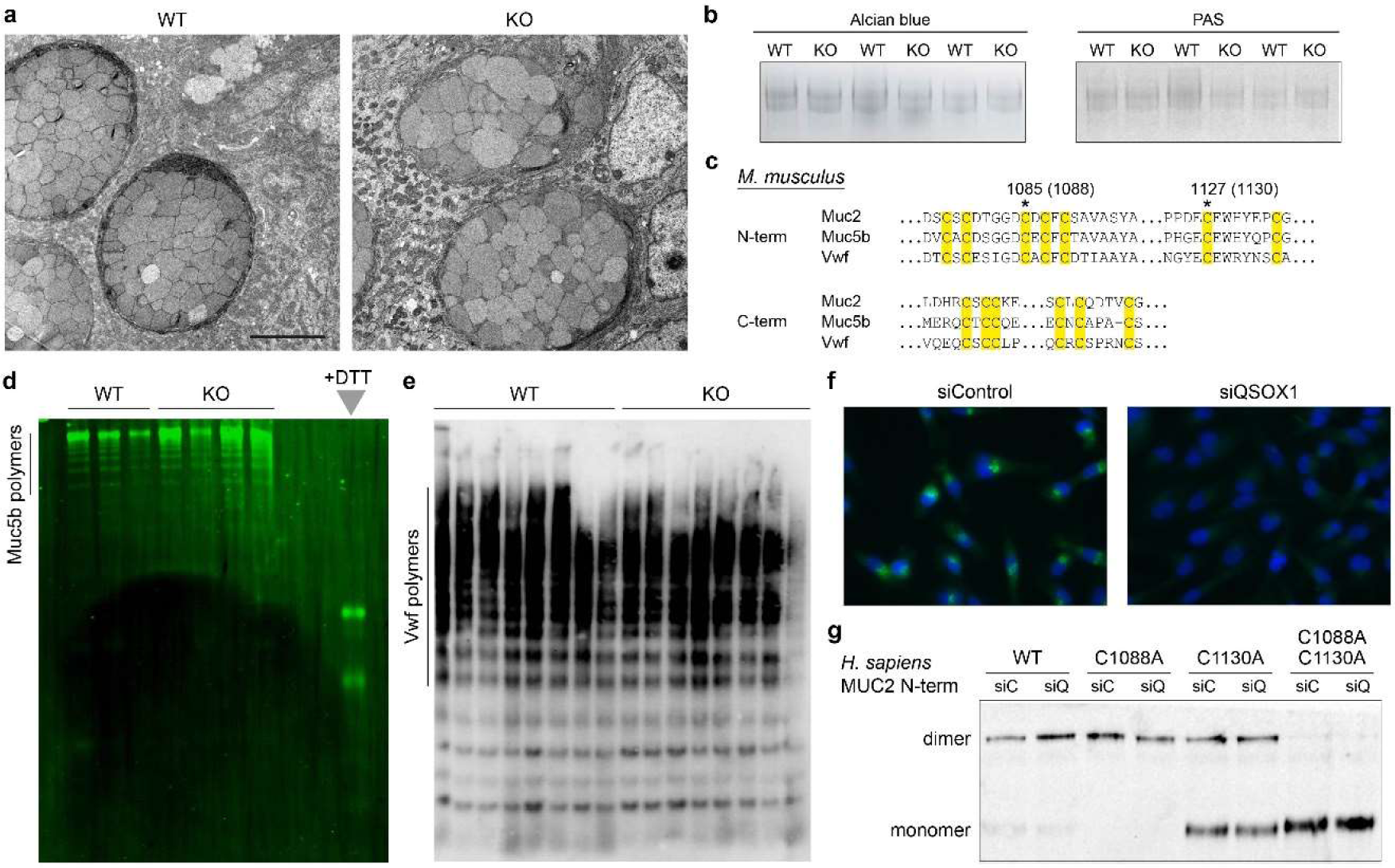
Normal mucin and VWF polymerization in QSOX1 KO mice. **a,** TEM micrographs show large and well-packed goblet cell granules in WT and KO colons. Scale bar is 5 µm. **b,** Alcian blue and PAS staining of reduced guanidine-insoluble mucins from WT and QSOX1 KO mice, separated on a 6% polyacrylamide gel. **c,** Amino acid sequences of murine Muc2 (Uniprot Q80Z19), Muc5b (Uniprot E9Q5I3), and Vwf (Uniprot Q8CIZ8) showing sequence conservation in the regions of the cysteines participating in intermolecular disulfide bonding to form polymers. Cysteines are highlighted in yellow, and asterisks indicate those that make intermolecular disulfides. Numbers indicate the amino acid positions of intermolecular disulfide-bonding cysteines in the amino-terminal region of murine Muc2, and in parentheses are corresponding amino acid positions in human MUC2, for reference to panel g. **d,** Muc5b fluorescent immunoblot of WT and QSOX1 KO lung lavage samples separated on agarose gels. **e,** Western blot analysis of blood Vwf from WT and QSOX1 KO mice. **f,** QSOX1 immunofluorescence (green) in control (siControl) and QSOX1 knockdown (siQSOX1) MDA-MB-231 cells. Blue is DAPI staining of nuclei. Scale bar is 20 µm. **g,** Western blot analysis of the MUC2 N-terminal region and indicated cysteine mutants in supernatants of transfected cell cultures. No differences in disulfide-mediated dimerization were observed for any MUC2 variant between siC (siControl) and siQ (siQSOX1).

### Mucins and related glycoproteins polymerize in QSOX1 knockout mice

Despite the normal appearance of goblet cell granules in the QSOX1 KO mice (Fig. 3a), it was possible that mucins were not properly polymerized prior to packaging and thus were unstable upon release into the intestinal lumen. Directly evaluating polymerization of colon mucins is technically difficult due to their excessive size and the insolubility of the assembled network^38^. We therefore took three complementary approaches to address this issue. First, we compared the guanidine-insoluble fraction of colon lysates, which contains Muc2, by staining with Alcian blue and periodic acid-Schiff (PAS). Using this protocol, we consistently observed similar amounts of staining for WT and KO, suggesting similar amounts of reducible polymerized mucins produced by goblet cells (Fig. 3b). Second, we took advantage of the fact that polymerized lung mucin Muc5b and the blood clotting glycoprotein von Willebrand factor (Vwf), which are homologous to Muc2 (Fig. 3c) and share the same polymerization mechanism^8, 39^, can be analyzed by agarose gel electrophoresis. Both Muc5b (Fig. 3d) and Vwf (Fig. 3e) from QSOX1 KO mice polymerized indistinguishably from their counterparts in WT animals. As a third approach to monitoring mucin disulfide bonding, we used a recombinant system comprising the amino-terminal ∼1400 amino-acid segment of human MUC2, which forms disulfide-bonded dimers that represent the crosslinking step proposed to occur in the Golgi^7, 8^. When this amino-terminal region of MUC2 was expressed in a cell line after depletion of QSOX1 (Fig. 3f), the same extent of dimerization occurred as in control cells, even after sensitization of the system by eliminating either of the two intermolecular disulfide crosslinks^7, 8^ (Fig. 3g). Extensive efforts to demonstrate differences in Muc2 polymerization by gels, electron microscopy, and atomic force microscopy (data not shown) did not yield evidence that mucin disulfide-mediated polymerization is impaired in the absence of QSOX1.

### Disulfide bonding in Golgi glycosyltransferases is perturbed in QSOX1 KO mice

In addition to disulfide-mediated crosslinking of mucins, another factor crucial to mucus function is glycosylation. Intriguingly, certain Golgi glycosyltransferases were reported to require disulfide bonds for function^40–42^. Specifically, mutation of cysteines in human sialyltransferase ST6GAL1 caused a decrease in functional heterodimer formation with partner glycosyltransferases and undermined the enhanced sialylation seen upon overexpression of wild-type ST6GAL1 in cells^41^. We therefore examined the redox states of Golgi sialyltransferases in QSOX1 KO mice. Lysates from isolated colon epithelial cells were treated with polyethylene-glycol modified maleimide (PEG-mal) of 2 kDa to alkylate reactive cysteine thiol groups that were not protected in disulfide bonds. Proteins were then separated by SDS-PAGE and western blotted for two sialyltransferases, St6gal1 and St3gal1, and for B3galt5, a galactosyltransferase that modifies core glycans^43^. In all three cases species that migrated more slowly, indicating the presence of cysteines conjugated to PEG-mal, were observed for QSOX1 KO samples (Fig. 4a-c). It should be noted that PEG conjugation retards protein migration to a greater extent than addition of the equivalent mass of protein^44^. As a control, treatment with the small (125 Da) cysteine alkylating agent N-ethylmaleimide (NEM) did not cause appreciable differences in glycosyltransferase migration (Fig. 4a-c). For comparison, no differences were noted between WT and KO in migration of the following proteins from PEG-mal-treated lysates: a representative ER oxidoreductase (Pdia4), a Golgi glycosyltransferase from the GALNT family (Galnt4), and the Golgi glycosyltransferase that generates the core 1 glycan (C1galt1) (Extended Data Fig. 4). These proteins have at least as many luminal cysteines as the glycosyltransferases analyzed in Figure 4.

**Fig. 4.**
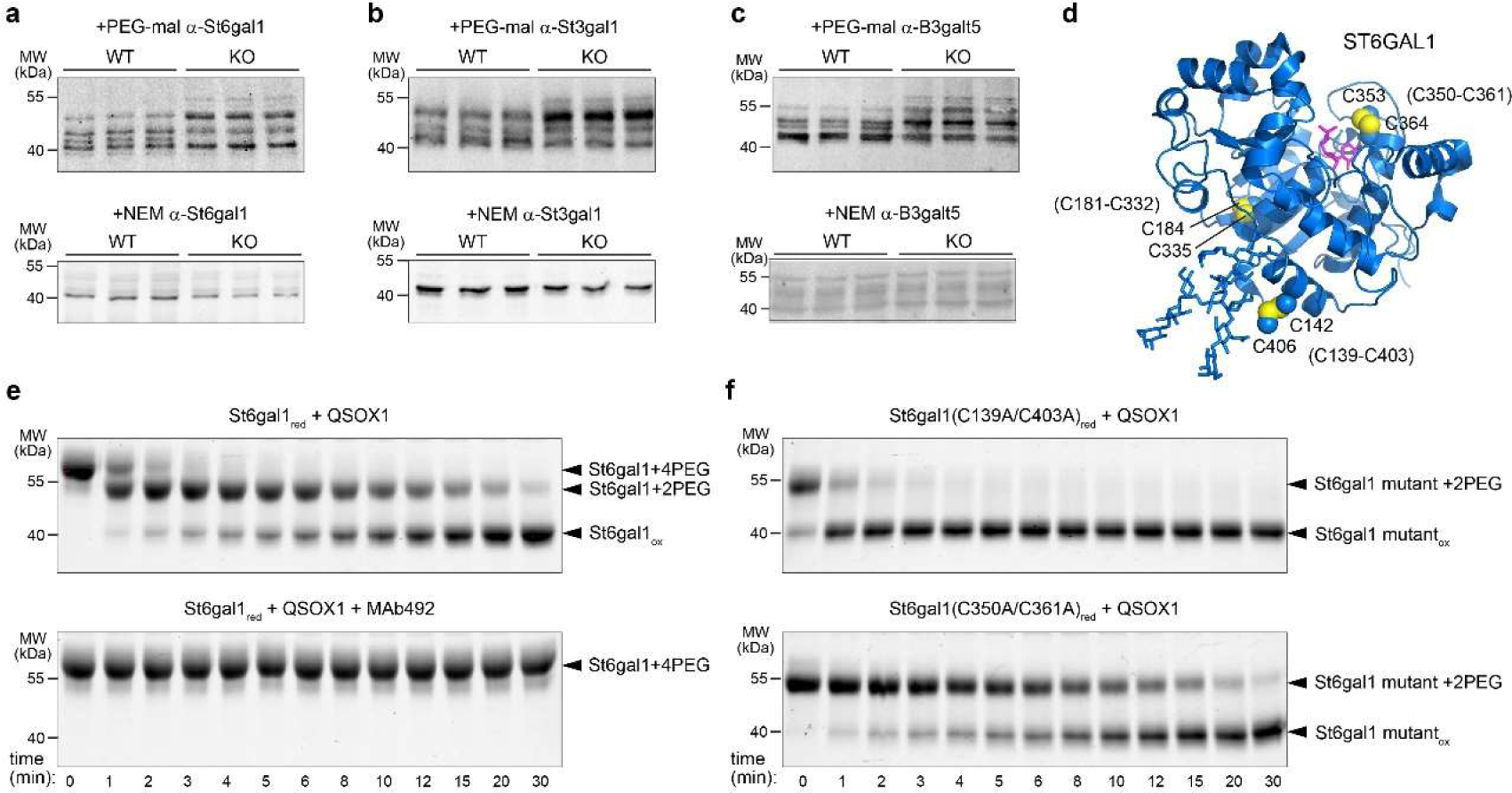
QSOX1 oxidizes Golgi glycosyltransferases. **a,** Western blot analysis of colon epithelial cell lysates from WT and QSOX1 KO mice. Lysates were treated with PEG-mal 2 kDa or NEM and probed using antibodies to St6gal1. **b,** As for panel a but using antibodies to St3gal1. **c**, As for panel a but using antibodies to B3galt5. **d**, Structure of the human ST6GAL1 catalytic domain in complex with CMP (PDB ID: 4JS2)^76^ with cysteine side chains shown as spheres and numbered. Numbering according to the murine St6gal1 sequence is indicated in parentheses. **e**, QSOX1 oxidizes St6gal1 in vitro. MAb492.1 is a monoclonal antibody that inhibits human QSOX1^73^. St6gal1 with one pair of free cysteines is modified by two PEG-mal additions (2PEG), whereas protein with two pairs of free cysteines gets four PEG-mal additions (4PEG). The change in migration per two PEG-mal modifications appears greater than 4 kD as expected due to the differences in hydrodynamic properties and SDS binding of PEG vs. protein^44^. **f,** QSOX1 oxidation of St6gal1 mutants. The C350-C361 disulfide (the redox-active disulfide present in the C139A-C403A mutant) is oxidized rapidly, while the C142-C406 disulfide (present in the C350A-C361A mutant) is oxidized slowly.

We next tested whether QSOX1 directly oxidizes Golgi sialyltransferases. Recombinant St6gal1 catalytic domain (Fig. 4d) was prepared, reduced, and incubated with purified QSOX1 for various times before addition of PEG-mal to quench oxidation reactions and label free thiols. St6gal1 was re-oxidized by QSOX1 as measured by resistance to PEG-mal modification (Fig. 4e). Notably, only two of the three St6gal1 disulfides in the recombinant protein (C142-C406 and C353-C364) were reactive in this experiment because the third (C184-C335) performs a stabilizing role within the structure^45^ (Fig. 4d) and is less accessible for reduction and modification. Of the two pairs of reduced cysteines, QSOX1 oxidized the pair adjacent to the St6gal1 active site much more rapidly than the pair involving the carboxy-terminal cysteine (Fig. 4e, f).

### Impaired glycosylation in colons of QSOX1 KO mice

The observed effect of QSOX1 on sialyltransferase redox state in vivo and in vitro would be functionally significant if manifested in altered glycosylation patterns. PAS staining is reported as a probe for sialic acid, as staining is decreased by prior treatment of colon sections with sialidase^46^. Though no major difference was detected in PAS in-gel staining of reduced guanidine-insoluble mucins (Fig. 3b), PAS staining of colon sections was much stronger for WT than KO (Fig. 5a). To further explore sialylation in QSOX1 KO colons, we used lectins that recognize relevant glycans (Fig. 5b). Sambucus nigra agglutinin (SNA) binds sialic acid preferentially in a α-2,6 linkage to terminal galactose^47^, which is the product of St6gal1 activity. Maackia amurensis lectin II (MAL II) binds sialic acid in a α-2,3 linkage^48^, the product of St3gal1 activity. Both SNA (Fig. 5c and Extended Data Fig. 5) and MAL II (Fig. 5d) labeled WT colons much more strongly than KO colons, indicating perturbed sialylation in QSOX1 KO animals. The sialylation defects were specific, since an antibody recognizing sialyl-Tn antigen (i.e., sialic acid on N-acetylgalactosamine (GalNAc) O-linked to serine or threonine), which is formed by the enzyme St6galnac1^49^ (Fig. 5f), labeled WT and KO colon tissue similarly (Extended Data Fig. 5).

**Fig. 5.**
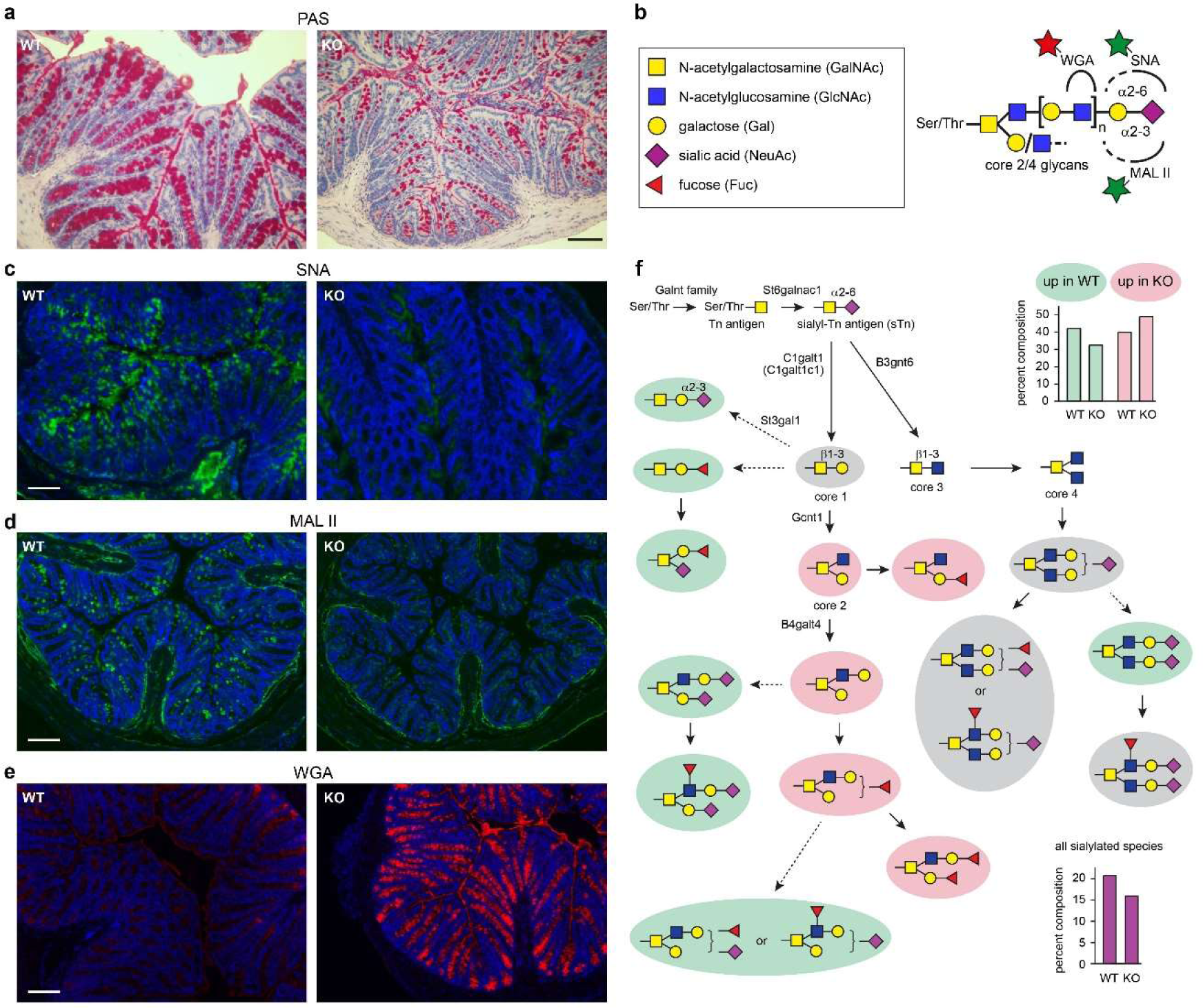
Altered glycan distribution in colons of QSOX1 KO mice. **a,** PAS and hematoxylin staining of colon cross sections. Scale bar is 50 µm. **b,** Schematic showing the glycan species recognized by the indicated lectins. Stars indicate fluorescent labels. **c,** Colon cross sections labeled with SNA lectin. Scale bar is 100 µm. **d,** Colon cross sections labeled with MAL II lectin. Scale bar is 100 µm. **e,** Colon cross sections labeled with WGA lectin. Scale bar is 100 µm. **f,** Pathway for O-glycan elaboration highlighted according to glycomics results. Paralogous enzymes also carry out some of the indicated steps. Gycans on a green background were populated in WT colon compared to QSOX1 KO colon. Glycans on a pink background were populated in the KO colon compared to the WT. Species on a gray background were similar in the two samples. Species with no background were not detected. Dashed arrows indicate decreased flux in the QSOX1 KO. The bar plot in the upper right shows the total percent composition of the glycan species with the corresponding colors. The bar plot in the lower right shows the total percent composition of all sialylated glycans detected.

Depressed sialylation would be expected to result in accumulation of precursor glycans. Specifically, GlcNAc would remain exposed without the coordinated action of galactosyl and sialyltransferases, which introduce the galactose and sialic acid terminal sugars (Fig. 5b). Indeed, wheat germ agglutinin (WGA), which recognizes GlcNAc (Fig. 5b), showed the opposite pattern to SNA and MAL II, labeling QSOX1 KO colons much more strongly than WT (Fig. 5e). Confirming that WGA labeling was due to GlcNAc binding rather than to other known activities of WGA^50^, a succinylated version of WGA that is more specific for GlcNAc^51^ showed similarly enhanced labeling of QSOX1 KO colons (Extended Fig. 5).

To confirm and extend the observations made with labeled lectins, we performed a comprehensive mass spectrometry glycomics analysis of colon epithelial extracts from WT and QSOX1 KO mice. This approach showed that O-linked glycans were less sialylated in the QSOX1 KO samples than in WT, and bottlenecks could be identified in glycosylation pathways (Fig. 5f). For example, sialylation of core 2 glycans was clearly compromised in the KO animals.

## DISCUSSION

The complex task of guarding the huge digestive and respiratory mucosal surfaces of the body requires a correspondingly complex macromolecular defense system. Thus, mucins have evolved diverse features to support their crucial protective functions. One is their ability to acquire intermolecular disulfide crosslinks and form polymers^52, 53^. Another feature is modification with thousands of O-linked glycans per mucin molecule^23^, facilitating mucus hydration and protecting the mucins against proteolytic degradation. Using these attributes, the multi-faceted mucin molecules form the physical and functional scaffold of the mucosal barrier. Our data reveal that the enzyme QSOX1 contributes to mucus hydrogel stability by activating Golgi sialyltransferases through disulfide bound formation, promoting expression of sialic acid in mucosal glycoproteins (Fig. 6).

**Fig. 6.**
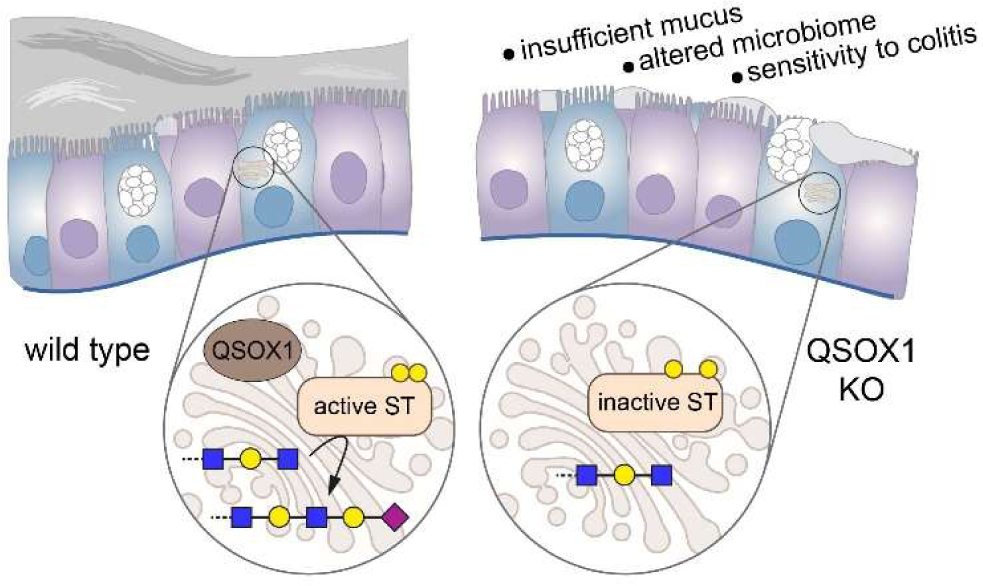
Role of Golgi QSOX1 and phenotypic effects of QSOX1 KO in the colon. Schematic diagram showing the effect of QSOX1 KO on sialylation in the Golgi and the impact on colon function. ST stands for sialyltransferase.

Based on our observation that colon mucus is compromised in QSOX1 KO mice, it was reasonable to speculate that disulfide bonding of mucins themselves would be defective, considering that mucins are thought to undergo disulfide-mediated polymerization in the Golgi apparatus^14^, where QSOX1 is localized^20, 54^. It was clear from the phenotype of the QSOX1 KO mice that earlier steps in mucin biosynthesis, i.e., those occurring in the ER, progressed normally, as these mice do not develop spontaneous colitis or rectal prolapse, which are phenotypes observed for a complete Muc2 KO^30^, Muc2 mutations that interfere with initial folding and assembly^55^, or KO of Agr2^18, 19^, which likely aids in early stages of Muc2 production. However, it was possible that a specific defect in mucin polymerization later in the secretory pathway led to the secretion of weak and poorly effective mucus, such that the QSOX1 KO mice had relatively normal gut function when unchallenged but were hypersensitive to induced colitis. Despite extensive investigation, however, we were not able to obtain evidence for a mucin disulfide-mediated polymerization defect.

It was previously observed that undermining O-glycosylation in the gut also leads to colitis. Mice lacking core 1 glycans, or both core 1 and core 3 glycans, exhibited spontaneous disease^56, 57^, whereas mice lacking only core 3 glycans were hypersensitive to induced colitis^58^. In contrast to the poor staining of mucins extracted from the mice unable to produce core glycans^57^, PAS and Alcian blue staining of the guanidine-insoluble fraction from QSOX1 KO colon extracts was apparently normal, and glycomics of colon epithelial cells confirmed core glycan production in the QSOX1 KO mice (Fig. 5f).

Building on the core O-glycans, colon mucins become highly elaborated by the action of additional glycosyltransferases adding glucose and galactose derivatives, and the glycan chain is finally capped by fucosylation and sialylation. Inspired by the observation that decreased disulfide bonding in certain sialyltransferases (STs) interferes with sialylation of N- and O-linked glycans in cultured cell lines^41^, we considered whether STs could be targets of QSOX1 activity in the colon. It is possible that the dependence of ST disulfides on oxygen tension reported by Hassinen et al.^41^ is due to the requirement for oxygen to drive disulfide formation by QSOX1^59^. By analyzing redox states in ex vivo colon epithelial cells, we found that cysteines of Golgi STs are indeed under-oxidized in the absence of QSOX1 (Fig. 4a-c). Moreover, we showed that the pathway linking disulfide formation to sialylation is critical for mucus stability and colon resistance to disease.

STs are a large family of enzymes, with 20 members in humans and mice^60, 61^. The size of the family likely represents the need for tissue-specific expression, temporal regulation, and specificity for distinct substrate sets. All ST family members retain a conserved pair of cysteines that participate in a structural disulfide bond^62^. However, the amino acid sequences, including the presence and position of other cysteine residues, have diverged quite substantially (Extended Data Fig. 6). Consequently, we do not expect that all members of the ST family will respond uniformly to QSOX1 activity. This diversity of potential redox handles offers the possibility of independently regulating sialylation of different substrates. Indeed, we observed that the levels of sialyl Tn antigen in WT and QSOX1 KO colons were similar, despite the different levels of other sialyl species. Interestingly, it was reported that sialylation can be tuned by differential localization of various STs to particular Golgi or post-Golgi compartments^63^. Interpreted together with the observations reported here, the types and extent of sialylation are likely to be a complex outcome of the levels of STs, their inherent capacity for redox-regulation, the physical and temporal extent of their co-localization with QSOX1, and the level of QSOX1 activity in the cell.

The importance of proper sialylation for mucus barrier integrity observed in our studies is reinforced by the finding that human ST mutations correlate with inflammatory bowel disease^64^. In a mouse model of a human mutation, deficiency in St6galnac1, a highly conserved ST that generates sialyl Tn antigen, resulted in phenotypes very closely resembling those of QSOX1 KO animals: a compromised mucus barrier, massive decrease in colon protein sialylation, changes in microbiome composition, and susceptibility to colitis^64^. In our experiments, antibody labeling of sialyl Tn showed comparable signal in WT and QSOX1 KO colons (Extended Data Fig. 5c,d), suggesting that St6galnac1 may not be a regulatory target of QSOX1. Nevertheless, the convergence of phenotypes in St6galnac1-defective and QSOX1 KO mice demonstrate that sialylation of multiple types is critical for gut health, and that STs can be perturbed both by mutation and by dysregulation.

QSOX1-mediated redox regulation of Golgi STs likely influences health and disease in contexts beyond the colon epithelium. For example, altered sialylation is a feature of cancer progression^65, 66^. In many cases, ST expression levels increase in cancer^67^, but it is possible that these enzymes are also activated post-translationally, since increased QSOX1 expression is seen in many cancer types^68–72^. Inhibition of extracellular QSOX1 using monoclonal antibody inhibitors^33, 73^ was found to slow tumor growth and decrease metastasis, most likely by modulating ECM properties rather than Golgi redox state^34^. Importantly, we show here that administration of QSOX1 inhibitory antibodies did not potentiate DSS-induced colitis (Fig. 1b), demonstrating that QSOX1 activity in colon physiology, and likely the Golgi functions of QSOX1 in general, are protected from inhibition by systemic inhibitory antibody treatment. Potential use of QSOX1 inhibitory antibodies as cancer therapy would thus not be expected to produce colitis as a side effect. It remains to be determined whether the increase in sialylation in cancer is linked to increased Golgi QSOX1, and whether this pathway would also be a valuable target for inhibition.

The key finding of this work is that the Golgi-localized and secreted enzyme QSOX1 contributes to intestinal physiology by regulation of Golgi glycosyltransferases rather than by direct introduction of disulfide bonds during mucin polymerization as hypothesized by us and others^74^. Our studies exposed an enzymatic, redox-controlled pathway for sialylation in the Golgi, which, due to the widespread expression of QSOX1^21^ and STs^60, 75^, is likely to play a role in other physiological processes in addition to the preservation of normal colon physiology and microbiome demonstrated here.

## Acknowledgements

The authors thank Diego Butera and Philip Hogg for confirming the observation that VWF multimers form in QSOX1 KO mice. Shalev Itzkovitz is gratefully acknowledged for helpful discussions. David Morgenstern validated St6gal1 mutants using mass spectrometry. EM studies were conducted at the Irving and Cherna Moskowitz Center for Nano and Bio-Nano Imaging at the Weizmann Institute of Science. Research was supported by the European Research Council under the European Union’s Seventh Framework Programme (ERC grant agreement 310649 to D.F.), the Mizutani Foundation for Glycoscience (to D.F.), the Israel Science Foundation (grant 2660/20 to D.F. and T.I.), the Center for Scientific Excellence at the Weizmann Institute of Science (to D.F.), the NIH (grants HL080396 and HL130938 to C.M.E.), the Department of Defense (grant W81XWH-17-1-0597), and the CF Foundation (JARAMI20F0 to A.M.J.H.). This research was also supported in part by the National Institutes of Health (NIH) grant R24GM137782-01 to P.A. at the Complex Carbohydrate Research Center.

## Author contributions

Conceptualization, T.I. and D.F.; Methodology, M.S., Y.E., O.B., N.V.; Investigation, T.I., N.R., P.A., N.M., N.Y., T.F., P.V., Z.L., A.M.J., C.M.E., D.F.; Writing – Original Draft, D.F.; Writing – Review & Editing, T.I. with contributions from all authors; Funding Acquisition, D.F., P.A., T.I., and C.M.E.

## Additional information

Correspondence and requests for materials should be addressed to D.F. or T.I.

## Extended Data Figures

**Extended Data Figure 1.**
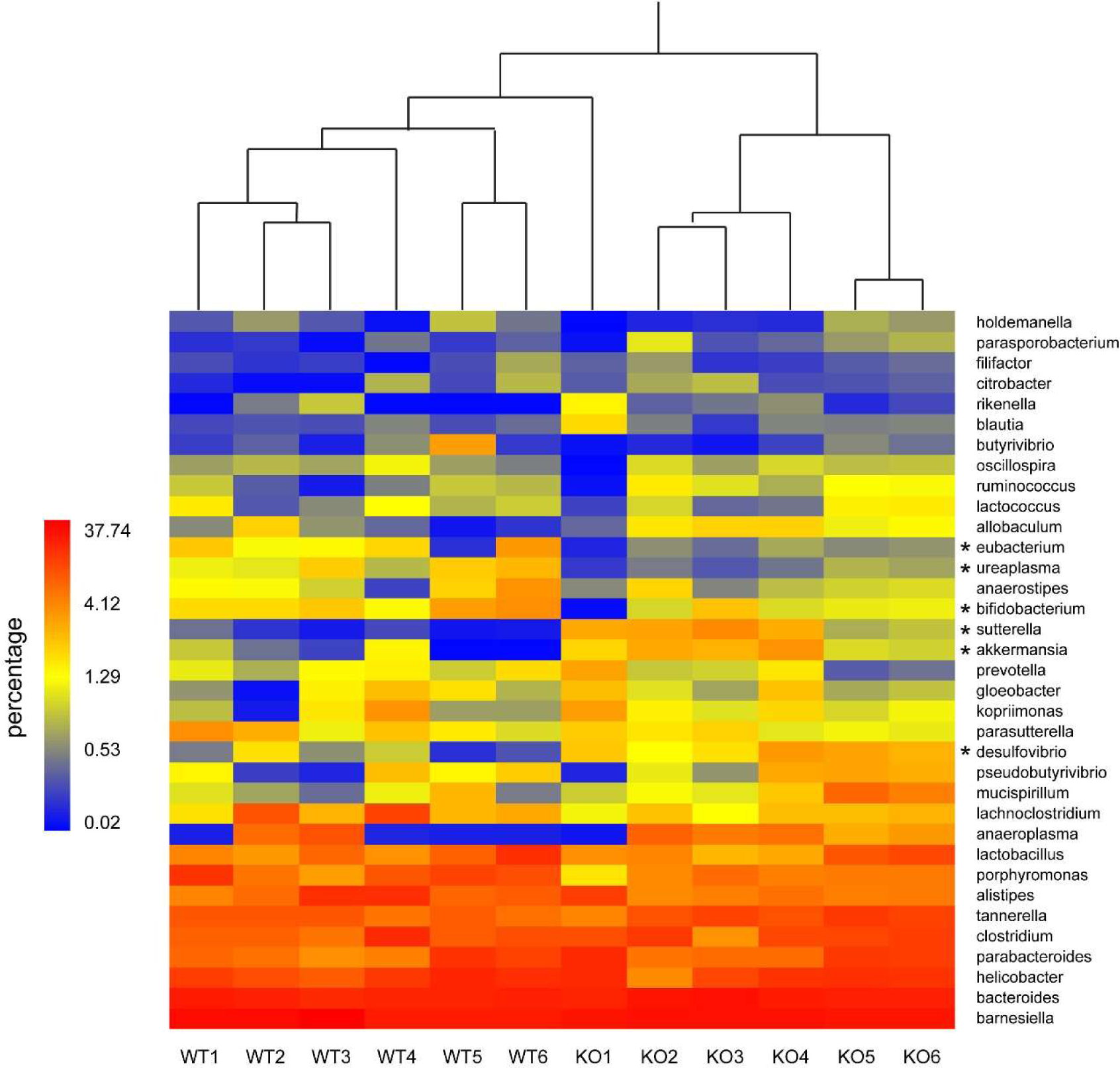
Microbial ecology of WT and QSOX1 KO mice. The predominant genera detected in the WT and QSOX1 KO fecal samples are displayed and clustered in a hierarchal dendrogram. The percentages of the genera are colored according to the scale on the left. The genera indicated by asterisks are plotted in Figure 2B. At the top of the heat map, the lengths of the connecting lines are related to the similarities between the microbial consortia of each sample. KO1 clusters loosely with the WT, but all other samples cluster according to genotype.

**Extended Data Figure 2.**
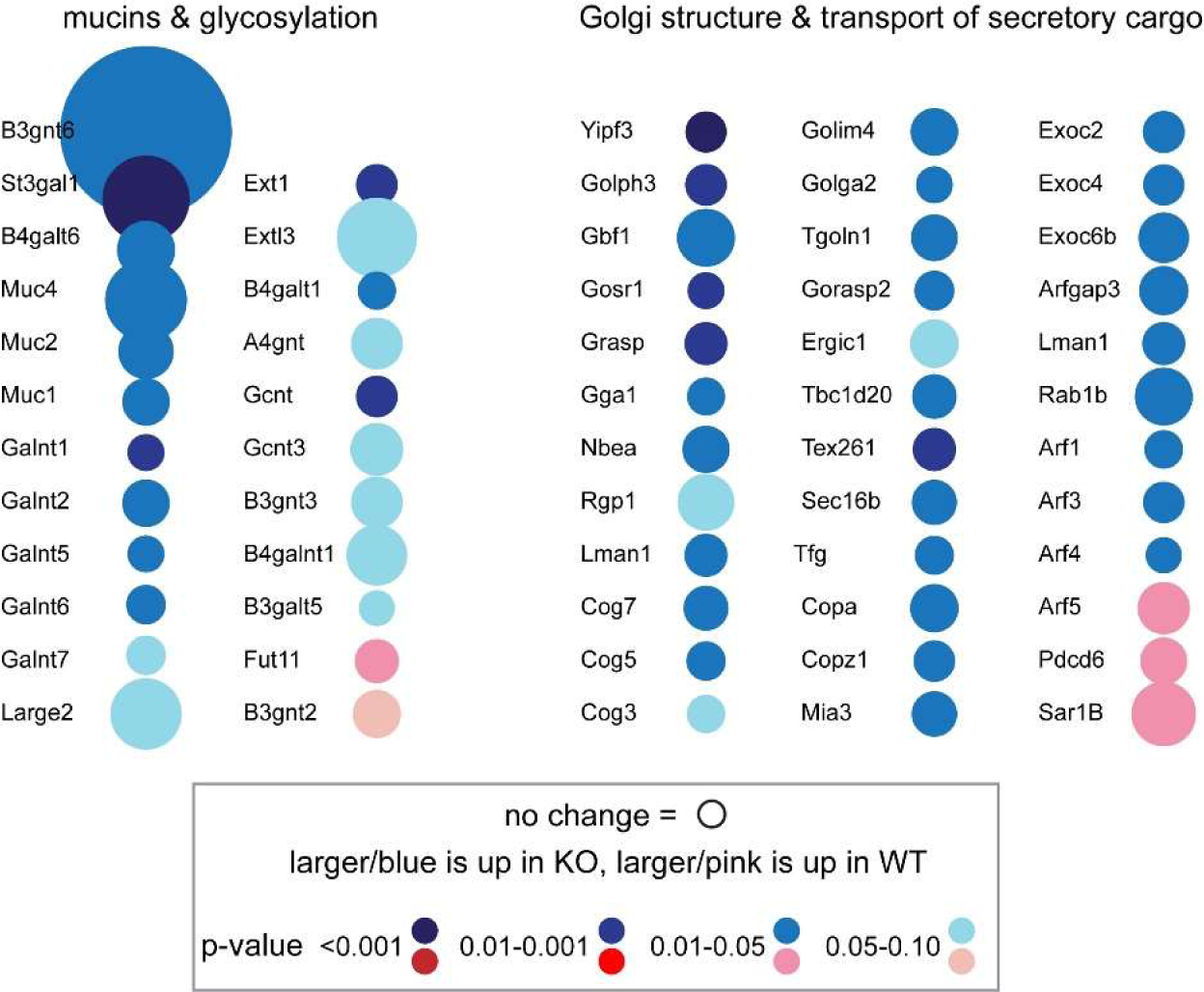
Summary of colon microarray. Changes in transcript levels of mucin genes and genes involved in glycosylation and vesicle transport in QSOX1 KO mice are displayed. The diameter of the circle is proportional to fold change, blue/pink is up/down-regulation in the QSOX1 KO relative to WT. Darker shades indicate lower p-values.

**Extended Data Figure 3.**
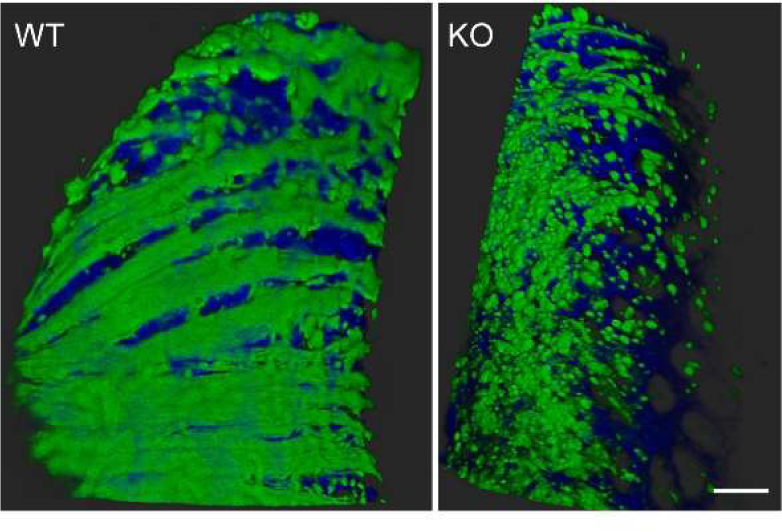
Additional images of colon longitudinal sections stained for Muc2. Even when regions of the WT and QSOX1 KO samples that have greater mucin coverage are selected, Muc2 in the KO does not appear as organized bundles. Scale bar is 50 µm.

**Extended Data Figure 4.**
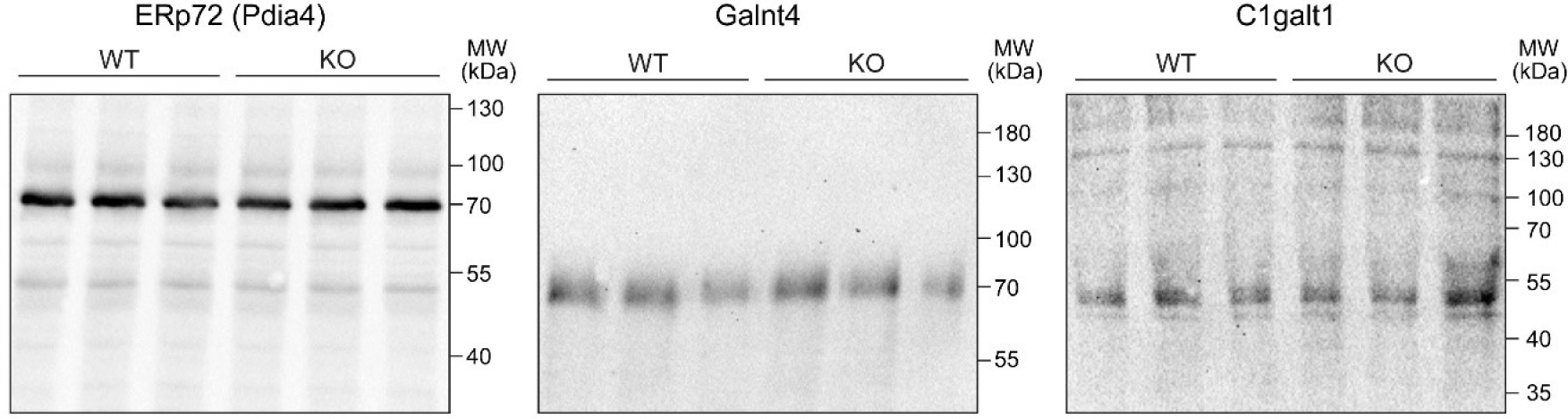
Western blot analysis of colon epithelial cell lysates from WT and QSOX1 KO mice. Lysates were treated with PEG-mal 2 kDa and probed using antibodies to ERp72, Galnt4, or C1galt1. No differences were seen in the migration of these proteins from QSOX1 KO mice compared to WT.

**Extended Data Figure 5.**
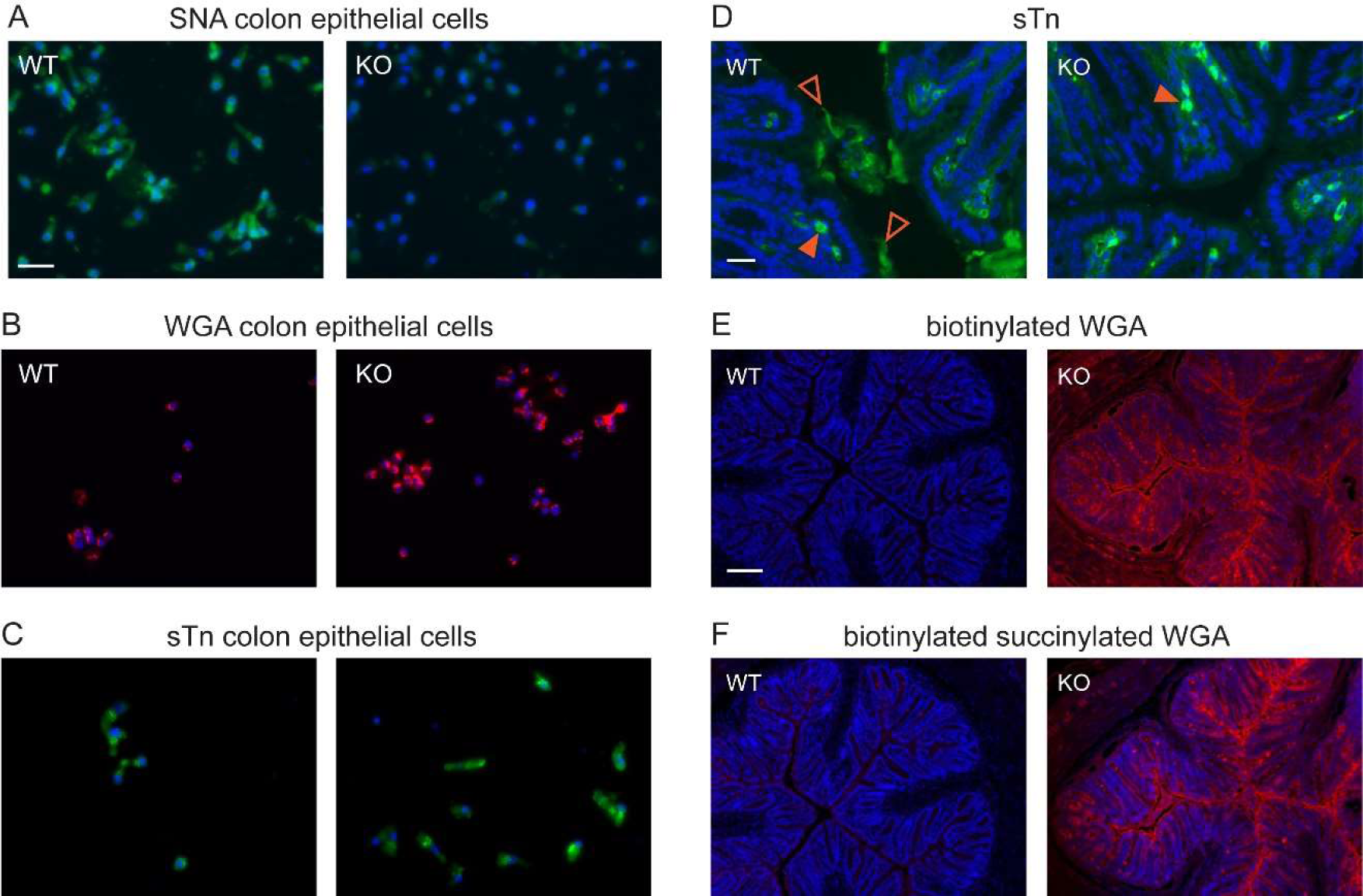
Additional lectin and antibody labeling of WT and QSOX1 KO colons and isolated epithelial cells. **a-c**, Isolated colon epithelial cells labeled with the indicated lectin (SNA or WGA) or antibody (α-sialyl Tn antigen (sTn)). Scale bars are 20 µm. **d**, Colon cross sections labeled with sTn antibody. Labeling was seen within the tissue in both WT and KO colons (filled arrowheads), but luminal staining was seen only in WT colons (open arrowheads) because little mucus was present in QSOX1 KO colon lumen. Scale bar is 20 µm. **e,f**, Succinylated and non-succinylated WGA both labeled KO colon sections much more strongly than WT colon sections. Scale bars are 100 µm.

**Extended Data Figure 6.**
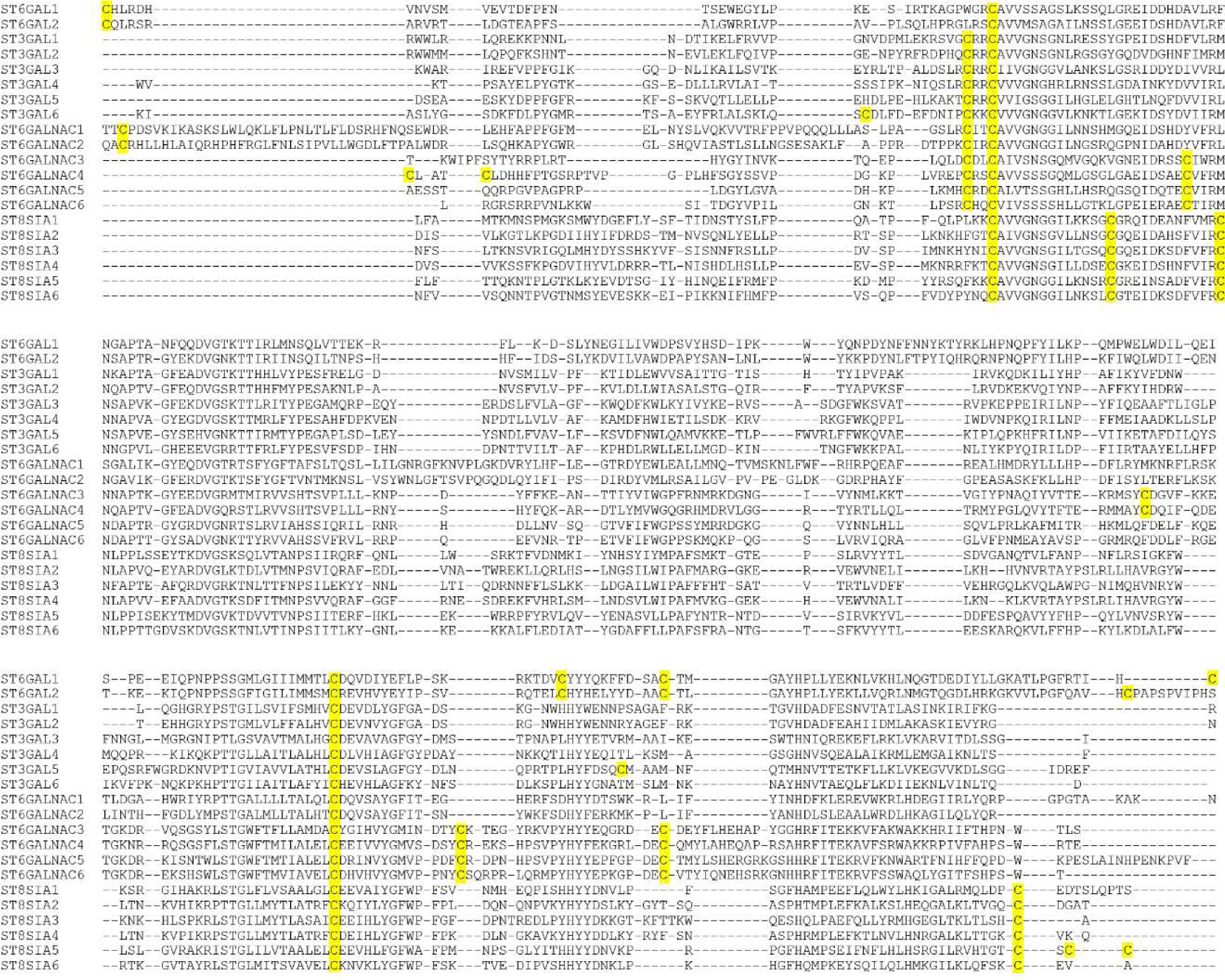
Sequence alignment of the catalytic domains of human sialyltransferases. Cysteine amino acids are highlighted. Two cysteines forming a structural disulfide are conserved in all sequences.

## Methods

### Generation of QSOX1 knockout mice

QSOX1 knockout (KO) mice were generated from the embryonic stem cell clone QSOX1^tm1a(KOMP)Wtsi^ obtained from KOMP Repository. The background strain is C57Bl. QSOX1 deletion was validated by mRNA and protein detection, and all mice were genotyped. WT and QSOX1 KO pairs in each of the experiments described in this work were littermates. All experiments were approved by the institutional animal care and use committee (IACUC) of the Weizmann Institute of Science. Approval numbers: 05480818-2, 01280121-2.

### Colon epithelial cell purification

The isolation protocol for colon epithelial cells was adopted from Bahar Halpern et al.^77^. Colons were removed, opened longitudinally, and incubated for 10 min in 5 ml of 10 mM ethylenediaminetetracetic acid (EDTA) in phosphate buffered saline (PBS) on ice. Colons were then moved to 5 ml of 100 µg/ml Liberase^TM^ (Sigma) in PBS and incubated for 2-3 hr at 37 °C with shaking. Finally, suspended material was filtered through a 100 µm cell strainer. Cells were washed in PBS and resuspended in RPMI-1640 medium supplemented with serum and antibiotics. For immunofluorescence and lectin labeling, cells were allowed to recover overnight (ON) at 37 °C with 5% CO2.

### Immunofluorescence and lectin labeling of cells

For immunofluorescence labeling, cells were transferred to glass cover slips in 24-well plates and after 1 hr at 37 °C were fixed for 20 min at room temperature (RT) with 3.7% formaldehyde. Following fixation, cells were permeabilized with 0.1% Triton X-100 in PBS for 2 min, washed, and incubated for 1 hr with 5% BSA in PBS containing 0.1% Tween (PBST). Cells were then incubated with primary antibody for 1 hr at RT, washed 3 times with PBST, and incubated with fluorescently-labeled secondary antibody and DAPI for 1 hr at RT. Following additional three washes, cover slips were placed, cells face down, onto a 5 µl drop of ProLong Gold antifade reagent (Invitrogen) on glass slides and were left to dry ON. For fluorescent lectin labeling (SNA-fluorescein and WGA-Alexa488), cells were fixed with formaldehyde, washed 3 times, and blocked with 3% BSA in PBS. Cells were then incubated with lectin (1:200 in PBST) and DAPI for 1-2 hr, washed 3 times, covered by mounting media, sealed with a cover slip, and allowed to dry ON prior to imaging. Biotinylated lectin labeling (WGA-biotin, succinylated WGA-biotin, and MAL II-biotin) was done similarly except an additional incubation with streptavidin-Alexa488, followed by washing, was performed.

### Imaging

All fluorescent light microscopy imaging, except for longitudinal sections, was done using an Olympus IX51 microscope equipped with Olympus XM10 camera. Images were analyzed using ImageJ software.

### Dextran sulfate sodium (DSS)-induced acute colitis

Acute colitis was induced by administration of DSS (2.5% in drinking water) for five days to 4 WT and 4 QSOX1 KO, 8-week-old female mice. Disease severity was assessed and scored on days 2, 4, and 5, as follows: 1= normal stool consistency, no rectal bleeding, 2= loose stool, no rectal bleeding, 3= diarrhea, minor rectal bleeding, and 4= gross rectal bleeding and prolapse. Mice were sacrificed at the experiment endpoint, colons were removed from cecum to rectum, and their lengths were measured. Distal colon was then separated and fixed for 2 hr in fresh, cold Carnoy’s solution (60% ethanol, 30% chloroform, 10% acetic acid) at 4 °C^78^ (Swidsinski et al., 2005). Following fixation, colons were washed in 100% ethanol and embedded in paraffin. Cross sections of distal colon were H&E stained, imaged, and analyzed. To test the potentiation of DSS-induced colitis by systemic QSOX1 inhibition, 30 mg/kg MAb316.1, an inhibitory monoclonal antibody for murine QSOX1^33^, was administered twice a week, for 3 weeks, via intraperitoneal injection to 5 WT 6-week-old female mice. A control group received control IgG in the same regimen. DSS was added to drinking water during the third week of antibody injection, and colitis was evaluated as described above.

### Microbiome analysis

Colons of 6 WT and 6 QSOX1 KO, 8-week-old female mice were removed, and feces (∼200 mg) closest to the rectum were collected into 200 µl lysis buffer (150 mM NaCl, 0.5% sodium deoxycholate, 0.1% Triton X-100, 50 mM Tris-HCl, pH 8.0, and protease inhibitors). Samples were lysed using Tissuelyzer LT bead mill (Qiagen), and DNA was purified using the MO BIO PowerSoil kit according to manufacturer’s instructions. Twenty µl of 50 ng/µl DNA of each sample were further processed for microbiome analysis by MR DNA (Molecular Research LP, Shallowater, TX, USA). Primers for the 16S rRNA gene V4 variable region 515/806 were used for amplification. Sequencing was performed on an Ion Torrent PGM following manufacturer’s guidelines. Sequence data were processed using a proprietary analysis pipeline. Barcodes, primers, and sequences less than 150 bp were removed. Sequences with ambiguous base calls and with homopolymer runs exceeding 6 bp were also removed. Sequences were denoised, operational taxonomic units (OTUs) were generated, and chimeras were removed. OTUs were defined by clustering at 3% divergence (97% similarity). Final OTUs were taxonomically classified using BLASTn against a database derived from RDPII and NCBI.

### Transcriptome microarray

Colons from 3 WT and 3 QSOX1 KO female mice were removed, and total RNA was immediately extracted using the RNeasy mini kit (Qiagen). RNA quality was assessed using the Bioanalyzer 2100 platform (Agilent), and samples were then processed and hybridized to GeneChip mouse gene 2.0 ST array according to the manufacturer’s instructions.

### Colon section preparation and labeling

With the exception of SNA labeling, immunofluorescence and lectin labeling of colons was done on paraffin sections. Colons were removed and immediately placed in fresh, cold Carnoy’s fixative solution for 2 hr at 4 °C, washed in 100% ethanol, and embedded in paraffin. For immunofluorescence labeling, sections were deparaffinized, and antigen retrieval was performed by incubation in 10 mM sodium citrate, pH 6, for 20 min at 95 °C. Slides were then washed in PBS, blocked in 3% BSA in PBS, and incubated with primary antibody (1:50-1:100 in PBST). Slides were then washed, incubated with fluorescently labeled secondary antibody (1:200 in PBST) and DAPI, and washed again. For both immunofluorescence and lectin labeling, sections were covered by mounting media, sealed with a cover slip, and allowed to dry ON prior to imaging. For WGA labeling, deparaffinized sections were blocked in 3% BSA in PBS and incubated with WGA-Alexa Fluor 488 (1:100 in PBST) and DAPI.

For MAL II labeling, deparaffinized sections were blocked in 3% BSA in PBS and incubated with MAL II-biotin (1:100 in PBST). Slides were then washed and incubated with streptavidin-Alexa Fluor 488 (1:200 in PBST) and DAPI. For S16 probe labeling, colon sections were deparaffinized, washed in PBS, and incubated with 100 µl of fluorescently labeled EUB (A488-GCTGCCTCCCGTAGGAGT) ON at 4 °C protected from light. Slides were then washed in PBST, incubated with DAPI for 1 hr, washed, covered by mounting media, and sealed with a cover slip.

For SNA staining in frozen sections, colons were removed, dissected into 2 mm sections, placed in cold tissue mold, covered with optimal cutting temperature (OCT) compound, immediately placed in liquid nitrogen, and then stored at −80 °C. For sectioning, frozen blocks were placed in a cryotome cryostat, and 10 µm thick sections were cut, placed on glass slides, and stored at −80 °C until labeling. Sections were fixed with 3.7% formaldehyde for 30 min at RT, washed 3 times in PBS, and incubated for 1 hr in 3% BSA in PBS. Sections were then incubated SNA-fluorescein (1:200 in PBST) and DAPI for 1 hr at RT, followed by washing 3 times with PBST. 5 µl of ProLong Gold antifade reagent (Invitrogen) was spread on a coverslip, which was then placed on the tissue section and left to dry ON prior to imaging.

### Longitudinal sectioning of colon and immunofluorescence labeling

Freshly removed colons were cut open longitudinally, gently washed with PBS containing Mg^2+^ and Ca^2+^, and covered by fresh, cold Carnoy’s fixative solution for 2 hr at 4 °C. Colons were then washed with PBS, incubated with primary antibody for 1 hr (1:50) in 0.1% PBST, washed again, and incubated with fluorescently labeled secondary antibody (1:200 in PBST) and DAPI. Following additional washing, the colons were covered by mounting media, sealed with a cover slip, and allowed to dry ON prior to imaging. Confocal imaging was performed using an upright Leica TCS SP8 (Leica microsystems CMS GmbH, Germany) at the Advanced Optical Imaging Unit, de Picciotto-Lesser Cell Observatory Unit of the Moross Integrated Cancer Center Life Science Core Facilities, Weizmann Institute of Science.

### Transmission electron microscopy

For preparation of thin cell sections, mouse colons were fixed using 2.5% gluteraldehyde, 2% paraformaldehyde in 0.1 M sodium cacodylate buffer, pH 7.4, at RT, washed in cacodylate buffer at 4°C, and post-fixed with 1% osmium tetroxide in cacodylate buffer for 1 hr, then incubated with 2% uranyl acetate for 1 hr. Samples were dehydrated in cold ethanol and then embedded in Epon. Thin sections were cut and stained with 2% uranyl acetate and Reynold’s lead citrate. Samples were visualized using a Tecnai T12 electron microscope (Thermo Fisher Scientific) equipped with a OneView camera (Gatan).

### Scanning electron microscopy (SEM)

Freshly removed colons were cut longitudinally, gently washed with PBS, and covered by fresh, cold Carnoy’s fixative solution for 1 hr at 4 °C. Following fixation, samples were gently rinsed in PBS and incubated in increasing concentrations of ethanol (30-100%). Samples in absolute ethanol were dried using critical point dehydration and stored under vacuum ON. Finally, samples were coated with 4 nm iridium and imaged using an ULTRA 55 FEG microscope (Zeiss).

### Insoluble mucin staining

Colons were removed, opened longitudinally, dissected into small pieces, and immediately placed in 0.5 ml cold lysis buffer containing 3 M guanidinium chloride. Colons were homogenized in a Tissuelyzer LT bead mill, and tissue debris was removed by 3 min centrifugation at 100g. The resulting supernatants were centrifuged for 10 min at 17000g, and the pellet was resuspended in 8 M urea dissolved in 20 mM Tris, pH 7.4, containing 50 mM tris(hydroxymethyl)phosphine (THP) and incubated at RT for 1 hr without agitation. The tube was inverted gently twice during this incubation. Remaining insoluble material was removed by centrifugation for 2 min at 17000g, and NEM was added to the supernatant to a concentration of 30 mM. After 1 hr at RT, aliquots were taken for separation on 1.5 mm thick 6% SDS-PAGE gels. After electrophoresis, gels were stained with Alcian blue and with PAS^79, 80^.

### Periodic Acid-Schiff (PAS) staining of colon sections

Paraffin colon sections were deparaffinized and immersed in Periodic acid solution for 5 min at RT followed by 3 washes with water. Slides were then immersed in Schiff’s reagent for 15 min at RT followed by washing in running tap water for 5 min. Finally, slides were counterstained in hematoxylin solution for 1 min, washed with running tap water, covered by mounting media, sealed with a cover slip, and allowed to dry ON prior to imaging.

### Western blots of Muc5b and Vwf polymers

For Muc5b analysis, lung lavage in saline was performed on WT and QSOX1 KO mice. Dot blots were performed to assess Muc5b levels in each lavage, and samples containing approximately equivalent amounts of Muc5b were separated on an SDS/agarose gel. After transfer of proteins from the gel to PVDF membrane by vacuum blot, the blot was probed with rabbit-anti-mouse Muc5b antibody^81, 82^.

For Vwf analysis, fresh blood samples were immediately mixed 1:10 with 3.2% sodium citrate pH 7.4, and plasma was collected after a 4000g spin for 20 min at RT. Plasma samples were diluted 1:20 and run on a 1% agarose gel prepared with TAE-SDS. Gel was transferred ON at 4 °C and 20 mA to methanol-activated PVDF membrane in Tris-glycine buffer containing 20% methanol. The membrane was probed with a rabbit-anti-VWF antibody.

### Analysis of MUC2 fragment dimerization

MDA-MB-231 cells were grown in 24-well plates. QSOX1-specific or scrambled siRNA oligonucleotides (20 nM) were transfected into the cells using jetPRIME reagent (Polyplus-transfection) according to the manufacturer’s instructions. Cells were allowed to recover for 1 day before plasmid DNA (0.5 µg in 500 µl) encoding the N-terminal segment of human MUC2^8^ was transfected into the same cells. Two days after siRNA transfection, culture medium was taken from transfected cell wells and analyzed by western blot. MUC2 was detected by polyclonal antibody raised against the MUC2 D3 region^8^. Validation of QSOX1 knockdown in this experiment was done by immunofluorescence using a polyclonal antibody against human QSOX1^20^.

### Colon epithelial cell lysis

Freshly isolated colon epithelial cells were pelleted and lysed by adding 10% trichloroacetic acid to the cell pellet and incubating on ice for 20 min.

Protein pellet was washed twice with cold acetone and dissolved in lysis buffer containing 10 mM of either PEG-mal 2 kDa or NEM. Thirty µg samples of cell lysate were separated on SDS-PAGE gels and western blotted for the indicated proteins.

St6gal1 preparation and oxidation assays. DNA coding sequence for amino acids 85 to 403 of murine St6gal1 was inserted downstream of the QSOX1 signal sequence (MRRCNSGSGPPPSLLLLLLWLLAVPGANA/AP, cleavage occurs at the slash), a His6 tag, and a TEV protease cleavage site in the pcDNA3.1 vector. Cysteine mutations were made in this plasmid using mutagenic PCR. The resulting plasmids were transiently transfected into HEK293F cells (ThermoFisher). Cells were maintained in FreeStyle 293 medium and transfected using the PEI Max reagent (Polysciences Inc.) with a 1:3 ratio (w/w) of DNA to PEI at a concentration of 1 million cells per ml. Six days after transfection, cells were removed from the cultures by centrifugation for 10 min at 500g. The culture media were then further clarified by centrifugation for 15 min at 9500g and filtration through a 0.45-μm pore-size membrane. St6ga11 and mutants were purified from the filtered medium by nickel-nitrilotriacetic acid (Ni-NTA) chromatography and concentrated to about 10 µM.

To prepare reduced St6gal1 and mutants, the proteins were incubated with 20 mM THP for 20 min at RT. Reduced St6gal1 and mutants were then applied to a PD-10 desalting column equilibrated with PBS plus 1 mM EDTA. The concentrations of the peak fractions were measured, and 200 µl reactions were set up containing 7 µM (WT) or 3.2 µM (mutants) reduced St6gal1 and initiated by addition of QSOX1 to a concentration of 100 nM. Aliquots (10 µl for WT and 12 µl for mutants) were removed from the reaction at the indicated times and added to 1 µl of 10 mM PEG-mal 2 kDa. Gel loading buffer was added, and aliquots were applied to SDS-PAGE prepared with 2,2,2-trichloroethanol. After electrophoretic separation, protein bands were visualized according to a stain-free protocol^83^.

O-glycome analysis. Protocols were based on published methods for glycan profiling^84^. Colon epithelial cells from 3 WT and 3 KO samples were homogenized in 1 ml high-salt buffer (2 M NaCl, 5 mM EDTA, 100 mM Tris, pH 7.4) by passing through a 23-gauge syringe needle 10 times before probe sonication for 2 min at 10 sec on/off intervals. The homogenate was then pelleted by centrifugation at 14000g for 30 min. The supernatant was discarded, and the pellet was resuspended in 1 ml urea lysis buffer (8 M urea, 200 mM Tris, pH 8.0, 100 mM DTT) and heated to 50 °C for 45 min to denature. Afterwards, the samples were alkylated by adding 55 mg iodoacetamide and incubating at room temperature for 45 min. The samples were desalted and buffer exchanged into 500 µl 50 mM ammonium bicarbonate with a 10-kDa molecular weight cutoff (MWCO) filter, after which 2 µL of Recombinant PNGaseF (Glycerol-free) (New England Biolabs, Ipswich, MA; catalog # P0709L) was added to each sample. The samples were incubated at 37 °C for 24 hr, with a second addition of 2 µl of PNGaseF after 24 hr for a total incubation time of 48 hr. Released N-glycans were removed using a 10-kDa MWCO filter by centrifuging for 15 min at 14000g, adding 400 µl 50 mM ammonium bicarbonate, and centrifuging again for 15 min at 14000g. The protein fraction containing O-linked glycans that remained in the filter was inverted into a clean microfuge tube and centrifuged. An additional 20 µl 50 mM ammonium bicarbonate was added to the filter and gently mixed by pipetting up and down before the filter was reinverted and centrifuged once more. The O-glycoprotein was then transferred to a clean glass vial that was set on dry ice to freeze and lyophilized overnight.

For β-elimination^85^, the lyophilized O-glycoproteins were dissolved in 250 µl 50 mM NaOH, after which 250 µl of 19 mg/250 µl NaBH4 in 50 mM NaOH was added, and the solution was heated at 45 °C for 18 hr. Samples were cooled and neutralized by adding 10% acetic acid dropwise. Samples were then desalted by passing through DOWEX H^+^ resin and C18 columns, lyophilized, and borates were removed using 10% acetic acid in methanol under a stream of N2. Released O-oligosaccharides were permethylated by using methyl iodide in a dimethyl sulfoxide (DMSO) and NaOH mixture^84^. The reaction was quenched with water, and the reaction mixture was extracted with dichloromethane and dried. The dried glycans were re-dissolved in methanol and profiled by matrix assisted laser desorption time of flight mass spectrometry (MALDI-TOF-MS). A 2 µl portion of each sample dissolved in methanol was added to 2 µl 2,5-dihydroxybenozic acid (DHB) matrix and spotted onto a MALDI plate. The samples were analyzed on an AB Sciex TOF/TOF 5800 System Mass Spectrometer using reflector positive ion mode. For electrospray ionization mass spectrometry (ESI-MS) analysis, dried permethylated O-glycans were dissolved in a solution of 100 µl water and 100 µl methanol containing 1 mM NaOH and were introduced into the mass spectrometer at a flow rate of 1 µL/min. Samples were analyzed on a Thermo Orbitrap Fusion Tribrid.

## References

1. Johansson, M. E. V., Sjövall, H. & Hansson, G. The gastrointestinal mucus system in health and disease. Nat. Rev. Gastroenterol. Hepatol. 10, 352–361 (2013).

2. Melhem, H., Regen-Komito, D. & Niess, J. H. (2021) Mucins dynamics in physiological and pathological conditions. Int. J. Mol. Sci. 22, 13642 (2021).

3. Wagner, C. E., Wheeler, K. M. & Ribbeck, K. Mucins and their role in shaping the functions of mucus barriers. Annu. Rev. Cell Dev. Biol. 34, 189–215 (2018).

4. Sharpe, C., Thornton, D. J. & Grencis, R. K. A sticky end for gastrointestinal helminths; the role of the mucus barrier. Parasite Immunol. 40, e12517 (2018).

5. Sicard, J.-F., Le Bihan, G., Vogeleer, P, Jacques, M. & Harel, J. Interactions of intestinal bacteria with components of the intestinal mucus. Front. Cell. Infect. Microbiol. 7, 387 (2017).

6. Braga Emidio, N., Brierley, S. M., Schroeder, C. I. & Muttenthaler, M. Structure, function, and therapeutic potential of the trefoil factor family in the gastrointestinal tract. ACS Pharmacol. Transl. Sci. 3, 583–597 (2020).

7. Javitt, G., Calvo, M. L. G., Albert, L., Reznik, N., Ilani, T., Diskin, R. & Fass, D. Intestinal gel forming mucins polymerize by disulfide-mediated dimerization of D3 domains. J. Mol. Biol. 431, 3740–3752 (2019).

8. Javitt, G., Khmelnitsky, L., Albert, L., Bigman, L. S., Elad, N., Morgenstern, D., Ilani, T., Levy, Y., Diskin, R. & Fass, D. Assembly mechanism of mucin and von Willebrand factor polymers. Cell, 183, 717–729 (2020).

9. Hughes, G. W., Ridley, C., Collins, R., Roseman, A., Ford, R. & Thornton, D. J. The MUC5B mucin polymer is dominated by repeating structural motifs and its topology is regulated by calcium and pH. Sci. Rep. 9, 17350. (2019)

10. Ridley, C., Lockhart-Cairns, M. P., Collins, R. F., Jowitt, T. A., Subramani, D. B., Kesimer, M., Baldock, C. & Thornton, D. J. The C-terminal dimerization domain of the respiratory mucin MUC5B functions in mucin stability and intracellular packaging before secretion. J. Biol. Chem. 294, 17105–17116 (2019).

11. Smillie, C. S. et al. Intra- and inter-cellular rewiring of the human colon during ulcerative colitis. Cell 178, 714–730 (2019).

12. Gouyer, V., Dubuquoy, L., Robbe-Masselot, C., Neut, C., Singer, E., Plet, S., Geboes, K., Desreumaux, P., Gottrand, F. & Desseyn, J. L. Delivery of a mucin domain enriched in cysteine residues strengthens the intestinal mucous barrier. Sci. Rep. 5, 9577 (2015).

13. Perez-Vilar, J., Eckhardt, A. E. & Hill, R. L. Porcine submaxillary mucin forms disulfide-bonded dimers between its carboxyl-terminal domains. J. Biol. Chem. 271, 9845–9850 (1996).

14. Perez-Vilar, J., Eckhardt, A. E., DeLuca, A. & Hill, R. L. Porcine submaxillary mucin forms disulfide-linked multimers through its amino-terminal D-domains. J. Biol. Chem. 273, 14442–14449 (1998).

15. Perez-Vilar, J. & Hill, R. L. The structure and assembly of secreted mucins. J. Biol. Chem. 274, 31751–31754 (1999).

16. Lippok, S., Kolšek, K., Löf, A., Eggert, D., Vanderlinden, W., Müller, J. P., König, G., Obser, T., Röhrs, K., Schneppenheim, S., Budde, U., Baldauf, C., Aponte-Santamaría, C., Gräter, F., Schneppenheim, R., Rädler, J. O. & Brehm, M. A. von Willebrand factor is dimerized by protein disulfide isomerase. Blood 127, 1183–1191 (2016).

17. Appenzeller-Herzog, C. & Ellgaard, L. The human PDI family: versatility packed into a single fold. Biochim. Biophys. Acta 1783, 535–548 (2008).

18. Park, S. W., Zhen, G., Verhaeghe, C., Nakagami, Y., Nguyenvu, L. T., Barczak, A. J., Killeen, N. & Erle, D. J. The protein disulfide isomerase Agr2 is essential for production of intestinal mucus. Proc. Natl. Acad. Sci. USA 106, 6950–6955 (2009).

19. Zhao, F., Edwards, R., Dizon, D., Afrasiabi, K., Mastroianni, J. R., Geyfman, M., Ouellette, A. J., Andersen, B. & Lipkin, S. M. Disruption of Paneth and goblet cell homeostasis and increased endoplasmic reticulum stress in Agr2^-/-^ mice. Dev. Biol. 338, 270–279 (2010).

20. Ilani, T., Alon, A., Grossman, I., Horowitz, B., Kartvelishvily, E., Cohen, S. R. & Fass, D. A secreted disulfide catalyst controls extracellular matrix composition and function. Science 341, 74–76 (2013).

21. Tury, A., Mairet-Coello, G., Esnard-Fève, A., Benayoun, B., Risold, P.-Y., Griffond, B. & Fellmann, D. Cell-specific localization of the sulphydryl oxidase QSOX in rat peripheral tissues. Cell Tissue Res. 323, 91–103 (2006).

22. Strous, G. J. & Dekker, J. Mucin-type glycoproteins. Crit. Rev. Biochem. Mol. Biol. 27, 57–92 (1992).

23. Arike, L. & Hansson, G. C. The densely O-glycosylated MUC2 mucin protects the intestine and provides food for the commensal bacteria. J. Mol. Biol. 428, 3221–3229 (2016).

24. Nyström, E. E. L., Martinez-Abad, B., Arike, L., Birchenough, G. M. H., Nonnecke, E. B., Castillo, P. A., Svensson, F., Bevins, C. L., Hansson, G. C. & Johansson, M. E. V. An intercrypt subpopulation of goblet cells is essential for colonic mucus barrier function. Science 372, eabb1590 (2021).

25. Carpenter, J., Wang, Y., Gupta, R., Li, Y., Prashamsha, H., Subramani, D.B., Reidel, B., Morton, L., Ridley, C., O’Neal, W. K., Buisine, M.-P., Ehre, C., Thornton, D. J. & Kesimer, M. Assembly and organization of the N-terminal region of mucin MU5AC: Indications for structural and functional distinction from MUC5B. Proc. Natl. Acad. Sci. USA 118, e2104490118 (2021).

26. Zanata, S. M., Luvizon, A. C., Batista, D. F., Ikegami, C. M., Pedrosa, F. O., Souza, E. M., Chaves, D. F. S., Caron, L. F., Pelizzari, J.V., Laurindo, F. R. M. & Nakao, L. S. High levels of active quiescin Q6 sulfhydryl oxidase (QSOX) are selectively present in fetal serum. Redox. Rep. 10, 319–323 (2005).

27. Portes, K. F., Ikegami, C. M., Getz, J., Martins, A. P., de Noronha, L., Zischler, L. F., Klassen, G., Camargo, A. A., Zanata, S. M., Bevilacqua, E. & Nakao, L. S. Tissue distribution of quiescin Q6/sulfhydryl oxidase (QSOX) in developing mouse. J. Mol. Hist. 39, 217–225 (2008).

28. Tabula Muris Consortium. Single-cell transcriptomics of 20 mouse organs creates a Tabula Muris. Nature 562, 367–372.

29. Caillard, A., Sadoune, M., Cescau, A., Meddour, M., Gandon, G., Polidano, E., Decayre, C., Da Silva, K., Manivet, P., Gomez, A.-M., et al. QSOX1, a novel actor of cardiac protection upon acute stress in mice. J. Mol. Cell. Cardiol. 119, 75–86 (2018).

30. Van der Sluis, M., De Koning, B. A. E., De Bruijn, A. C. J. M., Velcich, A., Meijerink, J. P. P., Van Goudoever, J. B., Büller, H. A., Dekker, J., Van Seuningen, I., Renes, I. B. & Einerhand, A. W. C. Muc2-deficient mice spontaneously develop colitis, indicating that Muc2 is critical for colonic protection. Gastroenterol. 131, 117–129 (2006).

31. Okayasu, I., Hatakeyama, S., Yamada, M., Ohkusa, T., Inagaki, Y. & Nakaya, R. A novel method in the induction of reliable experimental acute and chronic ulcerative colitis in mice. Gastroenterology 98, 694–702 (1990).

32. Eichele, D. D. & Kharbanda, K. K. (2017) Dextran sodium sulfate colitis murine model: an indispensable tool for advancing our understanding of inflammatory bowel disease pathogenesis. World J. Gastroenterol. 23, 6016–6029 (2017).

33. Grossman, I., Ilani, T., Fleishman, S. J. & Fass, D. Overcoming a species-specificity barrier in development of an inhibitory antibody targeting a modulator of tumor stroma. Protein Eng. Des. Sel. 29, 135–147 (2016).

34. Feldman, T., Grossman-Haham, I., Elkis, Y., Vilela, P., Moskovits, N., Barshack, I., Salame, T. M., Fass, D. & Ilani, T. Inhibition of fibroblast secreted QSOX1 perturbs extracellular matrix in the tumor microenvironment and decreases tumor growth and metastasis in murine cancer models. Oncotarget 11, 386–398 (2020).

35. Rooks, M. G., Veiga, P., Wardwell-Scott, L. H., Tickle, T., Segata, N., Michaud, M., Gallini, C. A., Beal, C., van Hylckama-Vlieg, J. E. T., Ballal, S. A., et al. Gut microbiome composition and function in experimental colitis during active disease and treatment-induced remission. ISME J. 8, 1403–1417 (2014).

36. Derrien, M., Vaughan, E. E., Plugge, C. M., and de Vos, W. M. Akkermansia municiphila gen. nov., sp. nov., a human intestinal mucin-degrading bacterium. Int. J. Syst. Evol. Microbiol. 54, 1469–1476 (2004).

37. Cani, P. D. & de Vos, W. M. Next-generation beneficial microbes: the case of Akkermansia muciniphila. Front. Microbiol. 8, 1765 (2017).

38. Herrmann, A., Davies, J. R., Lindell, G., Mårtensson, S., Packer, N. H., Swallow, D. M. & Carlstedt, I. Studies on the “insoluble” glycoprotein complex from human colon. Identification of reduction-insensitive MUC2 oligomers and C-terminal cleavage. J. Biol. Chem. 274, 15828–15836 (1999).

39. Dang, L. T., Purvis, A. R., Huang, R.-H., Westfield, L. A. & Sadler, J. E. Phylogenetic and functional analysis of histidine residues essential for pH-dependent multimerization of von Willebrand factor. J. Biol. Chem. 286, 25763–25769 (2011).

40. N. Hirano, Y., Suzuki, T., Matsumoto, T., Ishihara, Y., Takaki, Y., Kono, M., Dohmae, N. & Tsuji, S. (2012) Disulphide linkage in mouse ST6Gal-I: determination of linkage positions and mutant analysis. J. Biochem. 151, 197–203 (2012).

41. Hassinen, A., Khoder-Agha, F., Khosrowabadi, E., Mennerich, D., Harrus, D., Noel, M., Dimova, E. Y., Glumoff, T., Harduin-Lepers, A., Kietzmann, T. & Kellokumpu, S. A Golgi-associated redox switch regulates catalytic activation and cooperative functioning of ST6Gal-I with B4GalT-I. Redox Biol. 24, 101182 (2019).

42. Ortiz-Soto, M. E., Reising, S., Schlosser, A. & Seibel, J. Structural and functional role of disulphide bonds and substrate binding residues of the human beta-galactoside alpha-2,3-sialyltransferase 1 (hST3Gal1). Sci. Rep. 9, 17993 (2019).

43. Zhou, D., Berger, E. G. & Hennet, T. Molecular cloning of a human UDP-galactose:GlcNAcβ1,3GalNAc β1,3 galactosyltransferase gene encoding an O-linked core3-elongation enzyme. FEBS J. 263, 571–576 (1999).

44. Javitt, G., Cao, Z., Resnick, E., Gabizon, R., Bulleid, N. J. & Fass, D. Structure and electron-transfer pathway of the human methionine sulfoxide reductase MsrB3. Antioxid. Redox Signal. 33, 665–678 (2020).

45. Meng, L., Forouhor, F., Thieker, D., Gao, Z., Ramiah, A., Moniz, H., Xiang, Y., Seetharaman, J., Milaninia, S. & Su, M. Enzymatic basis for N-glycan sialylation: structure of rat α2,6-sialyltransferase (ST6GAL1) reveals conserved and unique features for glycan sialylation. J. Biol. Chem. 288, 34680–34698 (2013).

46. Roe, R., Corfield, A. P. & Williamson, C. N. Sialic acid in colonic mucin: an evaluation of modified PAS reactions in single and combination histochemical procedures. Histochemical J. 21, 216–222 (1989).

47. Shibuya, N., Goldstein, I. J., Broekaert, W. F., Nsimba-Lubaki, M., Peeters, B. & Peumans, W. J. The elderberry (Sambucus nigra L.) bark lectin recognizes the Neu5Ac(α2-6)Gal/GalNAc sequence. J. Biol. Chem. 262, 1596–1601 (1987).

48. Wang, W. C. & Cummings, R. D. The immobilized leukoagglutinin from the seeds of Maackia amurensis binds with high affinity to complex-type Asn-linked oligosaccharides. J. Biol. Chem. 263, 4576–4585 (1988).

49. Marcos, N. T., Bennett, E. P., Gomes, J., Magalhaes, A., Gomes, C., David, L., Dar, I., Jeanneau, C., DeFrees, S., Krustrup, D., Vogel, L. K., Kure, E. H., Burchell, J., Taylor-Papadimitriou, J., Clausen, H., Mandel, U. & Reis, C. A. ST6GalNAc-I controls expression of sialyl-Tn antigen in gastrointestinal tissues. Front. Biosci. 3,1443–5145 (2011).

50. Ryva, B., Zhang, K., Asthana, A., Wong, D., Vicioso, Y. & Parameswaran, R. Wheat germ agglutinin as a potential therapeutic agent for leukemia. Front. Oncol. 9, 100 (2019).

51. Monsigny, M., Roche, A. C., Sene, C., Maget-Dana, R. & Delmotte, F. Sugar-lectin interactions: how does wheat-germ agglutinin bind sialoglycoconjugates? Eur. J. Biochem. 104, 147–153 (1980).

52. Meldrum, O.W., Yakubov, G. E., Bonilla, M. R., Deshmukh, O., McGuckin, M. A. & Gidley, M. J. Mucin gel assembly is controlled by a collective action of non-mucin proteins, disulfide bridges, Ca2+-mediated links, and hydrogen bonding. Sci. Rep. 8, 5802 (2018).

53. Thornton, D. J., Sharpe, C. & Ridley, C. Intracellular processing of human secreted polymeric airway mucins. Ann. Am. Thorac. Soc. 15, S154–S158 (2018).

54. Tury, A., Mairet-Coello, G., Poncet, F., Jacquemard, C., Risold, P. Y., Fellmann, D. & Griffond, B. QSOX sulfhydryl oxidase in rat adenohypophysis: localization and regulation by estrogens. J. Endocrinol. 183, 353–363.

55. Heazlewood, C. K., Cook, M. C., Eri, R., Price, G. R., Tauro, S. B., Taupin, D., Thornton, D. J., Png, C. W., Crockford, T. L., Cornall, R. J., Adams, R., Kato, M., Nelms. K. A., Hong, N. A., Florin, T. H. J., Goodnow, C. C., and McGuckin, M. A. (2008) Aberrant mucin assembly in mice causes endoplasmic reticulum stress and spontaneous inflammation resembling ulcerative colitis. PLoS Med. 5, e54.

56. Fu, J., Wei, B., Wen, T., Johansson, M. E. V., Liu, X., Branford, E., Thomsson, K. A., McGee, S., Mansour, L., Tong, M., McDaniel, J. M., Sferra, T. J., Turner, J. R., Chen, H., Hansson, G. C., Braun, J. & Xia, L. Loss of intestinal core 1-derived O-glycans causes spontaneous colitis in mice. J. Clin. Invest. 121, 1657–1666 (2011).

57. Bergstrom, K., Fu, J., Johansson, M. E. V., Liu, X., Gao, N., Wu, Q., Song, J., McDaniel, J. M., McGee, S., Chen, W., Braun, J., Hansson, G. C. & Xia, L. Core 1- and 3-derived O-glycans collectively maintain the colonic mucus barrier and protect against spontaneous colitis in mice. Mucosal. Immunol. 10, 91–103 (2017).

58. An, G., Wei, B., Xia, B., McDaniel, J. M., Ju, T., Cummings, R. D., Braun, J. & Xia, L. Increased susceptibility to colitis and colorectal tumor in mice lacking core 3-derived O-glycans. J. Exp. Med. 204, 1417–1429 (2007).

59. Hoober, K. L., Joneja B, White, H. B., 3^rd^ & Thorpe, C. A sulfhydryl oxidase from chicken egg white. J. Biol. Chem. 271, 30510–30516 (1996).

60. Harduin-Lepers, A., Vallejo-Ruiz, V., Krzewinski-Recchi, M. A., Samyn-Petit, B., Julien, S. & Delannoy, P. The human sialyltransferase family. Biochimie 83, 727–737 (2001).

61. Lopez, P. H. H., Aja, S., Aoki, K., Seldin, M. M., Lei, X., Ronnett, G. V., Wong, G. W. & Schnaar, R. L. Mice lacking sialyltransferase ST3Gal-II develop late-onset obesity and insulin resistance. Glycobiol. 27, 129–139 (2017).

62. Rao, F. V., Rich, J. R., Rakić, B., Buddai, S., Schwartz, M. F., Johnson, K., Bowe, C., Wakarchuk, W. W., DeFrees, S., Withers, S. G. & Strynadka, N. C. J. Structural insight into mammalian sialyltransferases. Nat. Struct. Mol. Biol. 16, 1186–1188 (2009).

63. Kitano, M., Kizuka, Y., Sobajima, T., Nakano, M., Nakajima, K., Misaki, R., Itoyama, S., Harada, Y., Harada, A., Miyoshi, E. & Taniguchi, N. Rab11-mediated post-Golgi transport of the sialyltransferase ST3GAL4 suggests a new mechanism for regulating glycosylation. J. Biol. Chem. 296, 100354 (2021).

64. Yao, Y., et al. Mucus sialylation determines intestinal host-commensal homeostasis. Cell 185, 1172–1188 (2022).

65. Garnham, R., Scott, E., Livermore, K.E. & Munkley, J. ST6GAL1: A key player in cancer. Oncol. Lett. 18, 983–989 (2019).

66. Buffone, A. & Weaver, V. M. Don’t sugarcoat it: how glycocalyx composition influences cancer progression. J. Cell Biol. 219, e201910070 (2020).

67. Pietrobono, S. & Stecca, B. Aberrant sialylation in cancer: biomarker and potential target for therapeutic intervention? Cancers 13, 2014 (2021).

68. Antwi, K., Hostetter, G., Demeure, M. J., Katchman, B. A., Decker, G. A., Ruiz, Y., Sielaff, T. D., Koep, L. J. & Lake, D. F. Analysis of the plasma peptidome from pancreas cancer patients connects a peptide in plasma to overexpression of the parent protein in tumors. J. Protome Res. 8, 4722–4731 (2009).

69. Soloviev, M., Esteves, M. P., Amiri, F., Crompton, M. R. & Rider, C. C. Elevated transcription of the gene QSOX1 encoding quiescin Q6 sulfhydryl oxidase 1 in breast cancer. PLoS One 8, e57327 (2013).

70. Baek, J. A., Song, P. H., Ko, Y. & Gu, M. J. High expression of QSOX1 is associated with tumor invasiveness and high grades groups in prostate cancer. Path. Res. Pract. 214, 964–967 (2018).

71. Sung, H. J., Ahn, J. M., Yoon, Y. H., Na, S. S., Choi, Y. J., Kim, Y. I., Lee, S. Y., Lee, E. B., Cho, S. & Cho, J. Y. Quiescin Sulfhydryl Oxidase 1 (QSOX1) secreted by lung cancer cells promotes cancer metastasis. Int. J. Mol. Sci. 19, 3213 (2018).

72. Knutsvik, G., Collett, K., Arnes, J., Akslen, L. A. & Stefansson, I. M. QSOX1 expression is associated with aggressive tumor features and reduced survival in breast carcinomas. Mod. Pathol. 29, 1485–1491 (2016).

73. Grossman, I., Alon, A., Ilani, T. & Fass, D. An inhibitory antibody blocks the first step in the dithiol/disulfide relay mechanism of the enzyme QSOX1. J. Mol. Biol. 425, 4366–4378 (2013).

74. Dong, X. & Springer, T. A. Disulfide exchange in multimerization of von Willebrand factor and gel-forming mucins. Blood 137, 1263–1267 (2021).

75. Kitagawa H. & Paulson J. C. Differential expression of five sialyltransferase genes in human tissues. J. Biol. Chem. 269, 17872–17878 (1994).

76. Kuhn, B., Benz, J., Greif, M., Engel, A. M., Sobek, H. & Rudolph, M. G. The structure of human α-2,6-sialyltransferase reveals the binding mode of complex glycans. Acta Crystallogr. D. 69, 1826–1838 (2013).

77. Bahar Halpern, K., Massalha, H., Zwick, R. K., Moor, A. E., Castillo-Azofeifa, D., Rozenberg, M., Farack, L., Egozi, A., Miller, D. R., Averbukh, I., Harnik, Y., Weinberg-Corem, N., de Sauvage, F. J., Amit, I., Klein, O. D., Shoshkes-Carmel, M. & Itzkovitz, S. Lgr5+ telocytes are a signaling source at the intestinal villus tip. Nat. Commun. 11, 1936 (2020).

78. Swidsinski, A., Weber, J., Loening-Baucke, V., Hale, L. P. & Lochs, H. Spatial organization and composition of the mucosal flora in patients with inflammatory bowel disease. J. Clin. Microbiol. 43, 3380–3389 (2005).

79. Møller, H. J. & Poulssen, J. H. Staining of glycoproteins/proteoglycans in SDS-Gels. In The Protein Protocols Handbook, J. Walker, ed. (Humana Press), pp. 761–772 (2002).

80. Packer, N. H., Ball, M. S., Devine, P. L. & Patton, W. F. Detection of glycoproteins in gels and blots. In The Protein Protocols Handbook, J. Walker, ed. (Humana Press), pp. 761–772 (2002).

81. Zhu, Y., Ehre, C., Abdullah, L. H., Sheehan, J. K., Roy, M., Evans, C. M., Dickey, B. F. & Davis, C. W. Munc13-2^-/-^ baseline secretion defect reveals source of oligomeric mucins in mouse airways. J. Physiol. 586, 1977–1992 (2008).

82. Roy, M., et al. (2014) Muc5b is required for airway defense. Nature 505, 412–416 (2014).

83. Ladner, C. L., Yang, J., Turner, R. J. & Edwards, R. A. Visible fluorescent detection of proteins in polyacrylamide gels without staining. Anal. Biochem. 326, 13–20 (2004).

84. Shajahan, A., Supekar, N. T., Chapla, D., Heiss, C., Moremen, K. W. & Azadi, P. Simplifying glycan profiling through a high-throughput micropermethylation strategy. SLAS Technol. 25, 367–379 (2020).

85. Morelle W. & Michalski, J.-C. Analysis of protein glycosylation by mass spectrometry. Nat. Protoc. 2, 1585–1602 (2007).

